# Comparative Performance of Portable DNA Extraction Protocols and Bioinformatics Workflows for Rapid Detection of Pathogens and Antimicrobial Resistance Using Oxford Nanopore Sequencing

**DOI:** 10.64898/2026.02.23.707422

**Authors:** Louis Kyei-Tuffuor, Samuel Kekeli Agordzo, Angela Krobea Asante, Davina Serwaa Boateng, Winifred Nana Serwaa Adjei, Venus Nana Boakyewaa Frimpong, Rita Nartey Frimpong, Francis Agyemang-Yeboah, Rea Maja Kobialka, Ahmed Abd El Wahed, Uwe Truyen, Yaw Ampem Amoako, Richard Odame Phillips, Michael Frimpong

## Abstract

Oxford Nanopore technology (ONT) enables rapid, portable pathogen identification and detection of antimicrobial resistance (AMR). Still, the reliability of downstream genomic analyses is highly dependent on DNA extraction quality, particularly in resource-limited settings. This study comparatively evaluated four portable bacterial DNA extraction protocols, derived from three commercial kits, to determine their impact on nanopore sequencing performance, bioinformatics workflow completion, and field deployability. Six Gram-negative bacterial isolates (*Escherichia coli*, n= 4; *Pseudomonas* sp., n= 1; and *Salmonella* sp., n= 1) were processed using four extraction protocols: SwiftX DNA, SwiftX DNA with proteinase K (ProtK), SwiftX ParaBact, and NucleoSpin Microbial. Twenty-four DNA extracts (6 isolates x 4 protocols) were sequenced on a single multiplexed MinION R10.4.1 flow cell. Sequencing data were analysed using validated Galaxy-based generic and species-specific pipelines, with workflow completion defined as successful progression through quality control, assembly, virulence, plasmid and AMR detection modules. DNA purity varied substantially by extraction protocol and was strongly associated with workflow success. NucleoSpin Microbial achieved 100% workflow completion, SwiftX ParaBact achieved 83%, while both SwiftX DNA-based protocols failed to complete full workflows. Higher A260/A280 ratios were strongly correlated with successful workflow completion (Spearman’s ρ = 0.767, p < 0.0001). Importantly, key AMR genes required to classify isolates as multidrug resistant were consistently detected using both NucleoSpin Microbial and SwiftX ParaBact extractions. However, NucleoSpin Microbial assemblies showed significantly higher contiguity and enabled broader and more complete detection of virulence factors, pathogenicity islands, plasmid replicons, and accessory AMR genes, reflecting enhanced genomic resolution.

**IMPORTANCE:** Rapid whole-genome sequencing is increasingly used to detect antimicrobial resistance and guide public health responses, but its reliability depends strongly on how bacterial DNA is extracted. In this study, we show that DNA extraction method choice has a major impact on Oxford Nanopore sequencing performance across clinically relevant bacteria. While silica-column-based extraction maximised genomic completeness and analytical depth, paramagnetic bead-based reverse purification offered superior portability with sufficient resolution for frontline AMR surveillance. These findings highlight a practical trade-off between field deployability and high-resolution genomic characterisation in low-resource settings.

## 1. INTRODUCTION

Rapid and reliable identification of bacterial pathogens and their antibiotic resistance (ABR) determinants remains a major challenge for clinical laboratories, particularly in low- and middle-income countries (LMICs) where molecular diagnostic capacity is limited (Murray et al., 2022; Pokharel et al., 2019). Conventional culture-based methods typically require several days to yield results, delaying targeted antibacterial or antimicrobial therapy (Procop et al., 2020; Roach et al., 2025). Although Polymerase Chain Reaction (PCR)-based assays can shorten turnaround times, they rely on predefined targets and provide limited genomic context, thereby restricting their ability to capture the full spectrum of resistance and virulence profiles (Chin et al., 2021; Yang & Rothman, 2004). These limitations are especially pronounced in sub-Saharan Africa, where diagnostics for infectious diseases are largely centralised in a small number of well-resourced urban laboratories (Ofori et al., 2024). This model limits access to testing in peripheral facilities and contributes to major gaps in routine AMR surveillance, despite LMICs bearing a disproportionate burden of drug-resistant infections driven by high infectious disease prevalence, widespread and often unregulated antibiotic use, and uneven access to appropriate therapies (Elbehiry et al., 2025; Aruhomukama, 2022; Laxminarayan, 2022; Thornval et al., 2025).

Recognising this, the World Health Organisation identifies robust surveillance as a cornerstone of AMR control (W.H.O., 2014). While high-income countries benefit from established surveillance networks, many sub-Saharan African countries lack sustained national systems, resulting in fragmented or absent AMR data (Ogunleye et al., 2025; Seale et al., 2017). Addressing these gaps requires investment in laboratory capacity, data generation, and reporting infrastructure to support coordinated public health responses and evidence-based antimicrobial stewardship (Donkor et al., 2024).

Whole-genome sequencing (WGS) offers an unbiased approach for pathogen identification, enabling comprehensive detection of AMR genes, virulence factors, sequence types, and phylogenetic relationships from a single assay (Papamentzelopoulou et al., 2025; Yamin et al., 2023). However, widely used short-read platforms such as Illumina HiSeq and MiSeq, despite their accuracy, are constrained by large instrument footprints, complex workflows, and extended turnaround times, which limit their suitability for decentralised or field-based use (Hoenen et al., 2016; Liu et al., 2012). Portable, real-time Oxford Nanopore Technologies (ONT) devices, including the MinION, have therefore emerged as promising tools for rapid pathogen identification and AMR detection at or near the point of care, with several studies demonstrating culture-free bacterial diagnostics within hours (Lerminiaux et al., 2025; Oehler et al., 2025; Sakai et al., 2019). However, the performance of these nanopore-based workflows is tightly coupled to DNA extraction, particularly the recovery of long, intact fragments and the removal of inhibitory contaminants (Nishii et al., 2023; Oehler et al., 2025). DNA extraction methods optimised for qPCR or short-read sequencing may yield acceptable spectrophotometric purity and concentration yet perform poorly in nanopore sequencing, producing fragmented reads and low assembly contiguity that compromise downstream analyses (Kruasuwan et al., 2024; Purushothaman et al., 2024). This disconnect between bench-top quality metrics and real sequencing output is a major barrier to deploying rapid AMR diagnostics in the field, particularly in settings with limited resources (Papamentzelopoulou et al., 2025).

New “reverse purification” paramagnetic-bead extraction methods, such as the SwiftX kits, are specifically designed to keep DNA fragments long and intact even during fast, high-temperature lysis, unlike traditional silica-column protocols, which can easily break long DNA strands during the binding and elution steps (Byrne et al., 2022; Schurig et al., 2023, 2024). Whether these portable, paramagnetic bead-based protocols can match or outperform established silica-column kits for nanopore diagnostics in real-world field settings where portability, thermal robustness, and low per-sample costs matter most remains uncertain (Kuupiel et al., 2017; Pai et al., 2012; Schurig et al., 2024). To address this gap, the present study comparatively evaluates four bacterial DNA extraction protocols, assessing their impact on nanopore sequencing performance, bioinformatics workflow success, and practical field-deployability in resource-limited settings.

## 2.0 METHODOLOGY

### 2.1 Isolate selection and sample preparation

Six bacterial isolates, detailed in Table 1, were selected to evaluate the extraction protocols with Oxford Nanopore sequencing. The panel combined well-characterised reference strains and field isolates with clinically relevant resistance profiles. Two *Escherichia coli* strains from the European Reference Laboratory for Antimicrobial Resistance (EURL-AR) collection were included as external positive controls for third-generation cephalosporin and carbapenem resistance, providing stable standards for comparing extraction and sequencing performance.

Two additional *E. coli* isolates and one *Pseudomonas* sp. isolate recovered from poultry and cattle caecal contents as part of the ADAPT-KCCR One Health AMR surveillance study in Ghana were included to ensure that the evaluation covered diverse, field-derived Gram-negative pathogens with clinically relevant resistance profiles. These isolates were selected on MacConkey agar supplemented with cefotaxime or meropenem. An ATCC *Salmonella* sp. quality-control strain from the KCCR bacteriology unit was included as a routine laboratory reference, ensuring that protocol performance could be evaluated across key foodborne and opportunistic pathogens.

For all six isolates, stored cultures in Microbank cryovials or agar stabs were subcultured onto Columbia agar (Oxoid, United Kingdom) with 5% sheep blood and incubated at 37°C for 18 - 24 hours. Well-isolated colonies were subcultured to confirm purity and generate sufficient biomass for downstream DNA extraction and sequencing, in accordance with standard microbiological procedures (Sanders, 2012; Slattery et al., 2020)

**Table 1.**
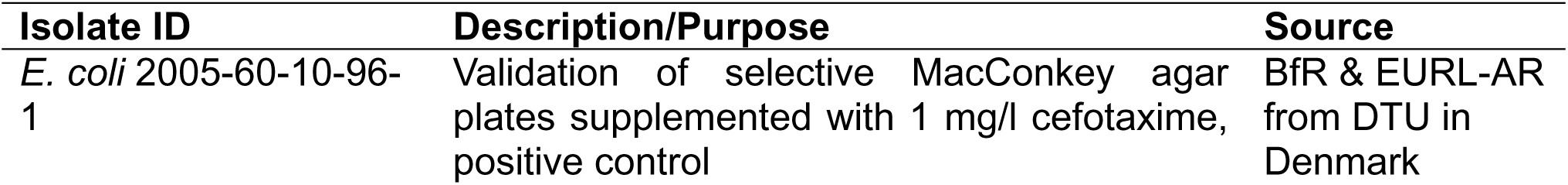
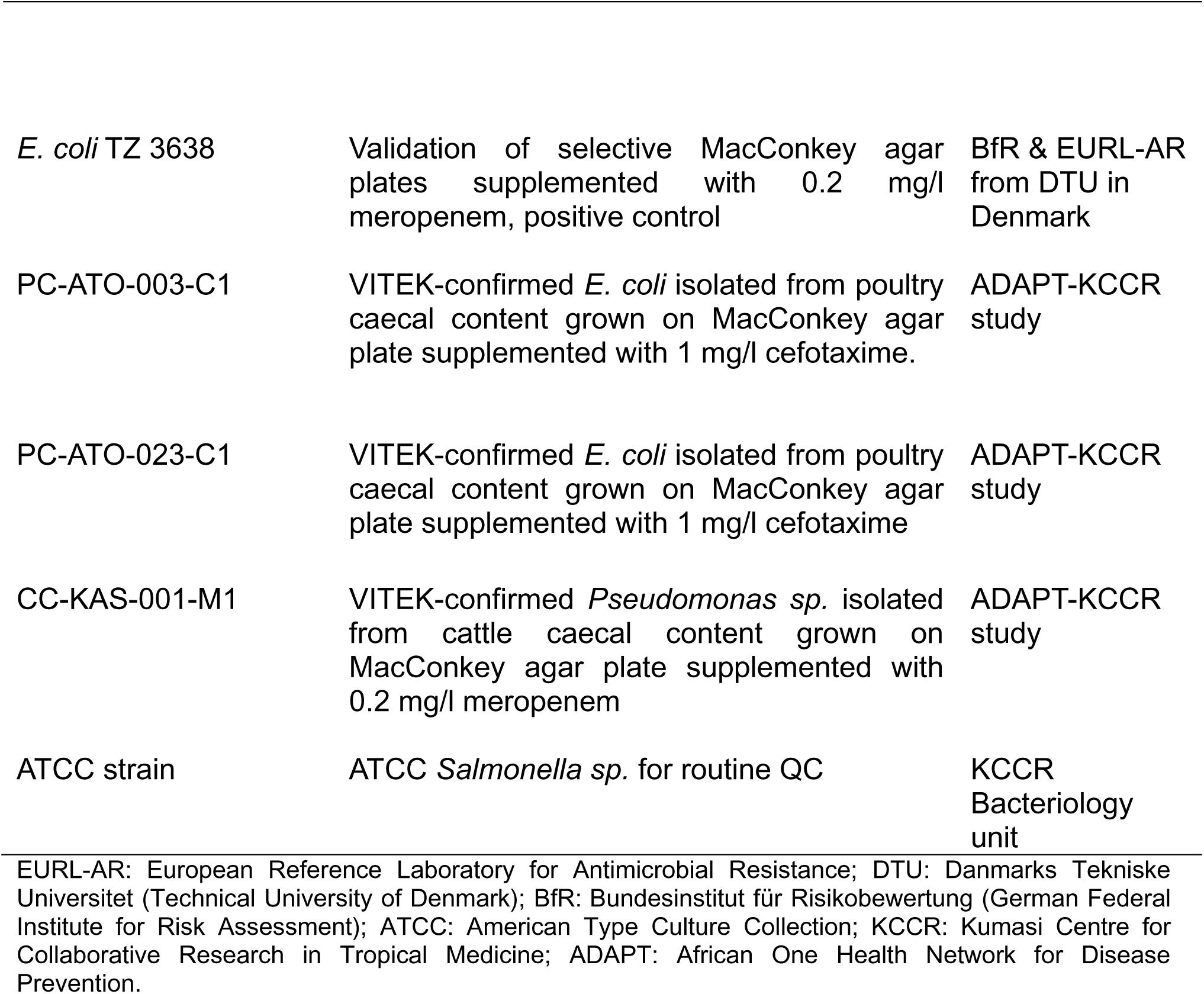
Bacterial Isolates used for the extraction protocol evaluation.

| Isolate ID | Description/Purpose | Source |
| --- | --- | --- |
| *E. coli* 2005-60-10-96-1 | Validation of selective MacConkey agar plates supplemented with 1 mg/l cefotaxime, positive control | BfR & EURL-AR from DTU in Denmark |
| *E. coli* TZ 3638 | Validation of selective MacConkey agar plates supplemented with 0.2 mg/l meropenem, positive control | BfR & EURL-AR from DTU in Denmark |
| PC-ATO-003-C1 | VITEK-confirmed *E. coli* isolated from poultry caecal content grown on MacConkey agar plate supplemented with 1 mg/l cefotaxime. | ADAPT-KCCR study |
| PC-ATO-023-C1 | VITEK-confirmed *E. coli* isolated from poultry caecal content grown on MacConkey agar plate supplemented with 1 mg/l cefotaxime | ADAPT-KCCR study |
| CC-KAS-001-M1 | VITEK-confirmed *Pseudomonas sp.* isolated from cattle caecal content grown on MacConkey agar plate supplemented with 0.2 mg/l meropenem | ADAPT-KCCR study |
| ATCC strain | ATCC *Salmonella sp.* for routine QC | KCCR Bacteriology unit |

EURL-AR: European Reference Laboratory for Antimicrobial Resistance; DTU: Danmarks Tekniske Universitet (Technical University of Denmark); BfR: Bundesinstitut für Risikobewertung (German Federal Institute for Risk Assessment); ATCC: American Type Culture Collection; KCCR: Kumasi Centre for Collaborative Research in Tropical Medicine; ADAPT: African One Health Network for Disease Prevention.

### 2.2 Genomic DNA (gDNA) extraction methods

To identify the most efficient protocol for bacterial DNA recovery, four extraction procedures based on three commercially available kits were evaluated across six diverse bacterial isolates. The protocols included that from the standard SwiftX DNA kit, a modified SwiftX DNA protocol incorporating Proteinase K digestion to enhance lysis of Gram-negative bacteria, the SwiftX ParaBact kit (Xpedite Diagnostics GmbH, Germany), and the NucleoSpin Microbial DNA kit (MACHEREY-NAGEL GmbH & Co. KG, Düren, Germany), which was used as the gold-standard comparator in this study. The kits spanned distinct lysis approaches (detergent, enzymatic, alkaline heat, and mechanical bead-beating) and purification methods (paramagnetic bead-mediated reverse purification and silica membrane spin-column). A schematic summary of the evaluated extraction procedures is shown in Figure 1.

For each isolate, genomic DNA was extracted from a single bacterial colony suspended in 700 µL of phosphate-buffered saline (Invitrogen PBS, Thermo Fisher Scientific). The SwiftX DNA protocol employed detergent-based lysis (Buffer DL) at 95°C, while the modified version incorporated a Proteinase K (ProtK) digestion step (60°C for 10 min) before detergent lysis to enhance recovery. The SwiftX ParaBact protocol utilised alkaline heat lysis (Buffer BPL) at 95°C. In contrast, the NucleoSpin Microbial protocol utilised mechanical disruption with glass beads and Proteinase K digestion, followed by column purification. In total, 24 gDNA extracts were generated (six isolates × four protocols), and each method was evaluated for DNA yield, purity, and suitability for downstream Nanopore sequencing, with particular attention to robustness and field adaptability. All extractions followed the manufacturers’ instructions, with minor optimisations as detailed in Supplementary File S1.

**Figure 1.**
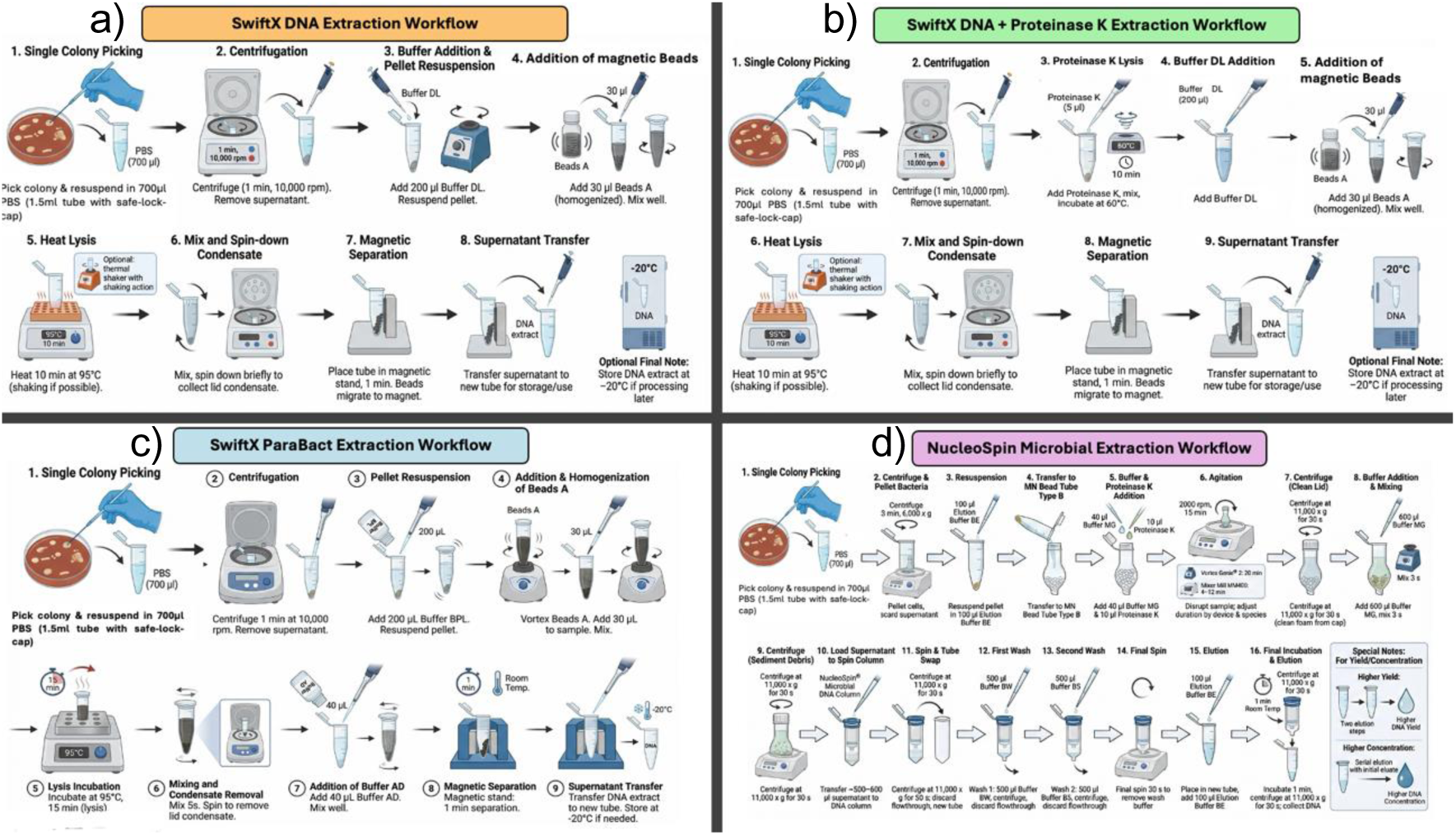
Schematic comparison of the four DNA extraction workflows evaluated. **(a)** SwiftX DNA: Detergent-based lysis with reverse purification. **(b)** SwiftX DNA + ProtK: Modified protocol with Proteinase K digestion. **(c)** SwiftX ParaBact: Alkaline heat lysis with reverse purification. (**d)** NucleoSpin Microbial: Mechanical bead-beating and enzymatic lysis followed by silica-column purification.

### 2.3 DNA quality assessment and quantification

DNA purity was assessed by microvolume UV spectrophotometry using a DeNovix DS-11 instrument (DeNovix Inc., Wilmington, USA). For each extract, 1.0 µL of DNA was applied to the lower pedestal, the arm was lowered to form a liquid column, and absorbance at 260 nm and 280 nm was measured in triplicate. A260/A280 ratios were recorded, and mean values were used for analysis; ratios between 1.8 and 2.0 were interpreted as indicative of acceptable DNA purity. DNA concentration was determined fluorometrically using a Qubit 4 Fluorometer with the Qubit dsDNA High Sensitivity (HS) Assay Kit (Invitrogen, Thermo Fisher Scientific Inc., USA). The fluorometer was calibrated with the two supplied standards, and a working solution was prepared according to the manufacturer’s instructions. For each sample, 2 µL of DNA was mixed with 198 µL of working solution in a Qubit assay tube, briefly vortexed, and incubated at room temperature for 2 minutes before reading. Measurements were performed in triplicate, and the instrument reported DNA concentration in ng/µL based on the standard curve.

### 2.4 Whole genome sequencing (WGS) of isolates

Whole-genome sequencing of the 24 DNA extracts was performed on the Oxford Nanopore MinION platform using the Rapid Barcoding Kit 24 V14 (SQK-RBK114.24; Oxford Nanopore Technologies, Oxford, UK), following the Nanopore-Only Microbial Isolate Sequencing Solution (NO-MISS) workflow. All 24 DNA samples were multiplexed and sequenced on a single MinION flow cell, enabling direct comparison of extraction performance under identical sequencing conditions.

For library preparation, 200 ng of purified genomic DNA per sample was used. DNA input was confirmed using a Qubit 4 Fluorometer, and 200 ng in 10 µL was combined with a unique Rapid Barcode (RB01–RB24) and subjected to the standard 30 °C/80 °C tagmentation program, according to the manufacturer’s instructions. Barcoded samples were pooled and purified using a 1:1 AMPure XP bead clean-up, washed with 80% ethanol, and eluted iElution Buffer to generate a single multiplexed library. Sequencing was performed on an R10.4.1 flow cell (FLO-MIN114) mounted on a MinION Mk1B device. Flow cell priming and library loading were carried out according to the manufacturer’s protocol. The sequencing library was prepared by combining Sequencing Buffer, Library Beads, and 12 µL of the pooled DNA library, and loaded via the SpotON sample port.

The sequencing was controlled using MinKNOW software v25.05. The run time was set to 72 hours with a minimum read length of 200 bp. High-accuracy (HAC) real-time basecalling and barcode demultiplexing (RB01–RB24) were enabled, generating POD5 and FASTQ outputs. The sequencing configuration targeted ≥50× genome coverage per sample, sufficient for downstream genome assembly, plasmid reconstruction, antimicrobial resistance profiling, and cgMLST/wgMLST analyses.

### 2.5 Preliminary Galaxy-based generic analysis

Bioinformatic pre-processing and initial analysis were performed for all 24 Nanopore-sequenced samples on Galaxy.eu, adapting the gene-based pathogen identification workflow from the tutorial “Pathogen detection from (direct Nanopore) sequencing data using Galaxy-Foodborne Edition” (Batut et al., 2025; Hiltemann et al., 2023). This preliminary analysis was conducted before downstream species-specific bioinformatics pipelines and served to evaluate data quality, confirm taxonomy, and obtain first-pass genomic feature annotations.

Raw FASTQ reads were subjected to quality assessment using Falco v1.2.4 (Smith & de Sena Brandine, 2021) and NanoPlot v1.46.1 (De Coster et al., 2018). Adapter trimming was performed with Porechop v0.2.4 (Wick et al., 2017), and additional read filtering (minimum length and quality) was performed with fastp v1.0.1(Chen et al., 2018). Summary reports from all pre-processing steps were collated with MultiQC v1.27 (Ewels et al., 2016). Taxonomic profiling was conducted using Kraken2 v2.1.3 (Wood & Salzberg, 2014) against both the Standard+PF and Kalamari databases (Katz et al., 2025). High-quality reads were assembled de novo with Flye v2.9.6 (Kolmogorov et al., 2019), and assemblies were polished to consensus with Medaka v2.1.1(Oxford Nanopore Technologies, 2023; Lamas et al., 2023).

Virulence genes were predicted with ABRicate 1.0.1 using the VFDB database (Chen et al., 2004), and antimicrobial resistance genes were detected with AMRFinderPlus v3.12.8 (Feldgarden et al., 2019) against the NCBI AMR database. Putative plasmid replicons were identified by screening assemblies with PlasmidFinder v2.1.6 (Carattoli et al., 2014; Carattoli & Hasman, 2020) against a plasmid replicon database, and all tabular outputs were exported from Galaxy for downstream comparison of extraction protocols across isolates.

### 2.6 Species-specific bioinformatics analysis

For the four *E. coli* isolates, species-level characterisation was performed with the Sciensano STEC Pipeline v1.2 on the Galaxy External instance, which has been validated for routine WGS-based STEC typing (Bogaerts et al., 2021; Galaxy External, 2025; Nouws et al., 2023). This “push-button” workflow carries out read trimming and basic quality control (Chen, 2023; Shen et al., 2024), de novo assembly (Bankevich et al., 2012; Gurevich et al., 2013; Kolmogorov et al., 2019), and automated species confirmation (Li, 2021; Li et al., 2009; Low et al., 2019), followed by integrated assays for antimicrobial resistance gene detection (Bortolaia et al., 2020; Feldgarden et al., 2019), virulence gene characterization (Joensen et al., 2014), O: H serotype prediction (Joensen et al., 2015), plasmid replicon identification (Carattoli & Hasman, 2020), and sequence typing using MLST/cgMLST schemes linked to curated public databases such as EnteroBase and PubMLST (Abueg et al., 2024; Bogaerts et al., 2025; Jaureguy et al., 2008; Zhou et al., 2020).

The ATCC *Salmonella* quality-control isolates were analysed with the Sciensano Salmonella Pipeline v1.0 on Galaxy External (Bogaerts et al., 2025), which mirrors the structure of the STEC workflow but implements Salmonella-specific assays. After read trimming, quality control, and de novo assembly, the pipeline performs species confirmation, in silico serotyping (e.g., using SISTR v1.1.1) (Yoshida et al., 2016; Zhang et al., 2019), antimicrobial resistance gene detection (Bortolaia et al., 2020; Feldgarden et al., 2019), prediction of *Salmonella* pathogenicity islands (Kombade et al., 2021; Siriken, 2013), virulence gene profiling (Jin et al., 2024), and plasmid replicon detection (Robertson & Nash, 2018), again drawing on continuously synchronised reference databases to support public health surveillance applications (Inouye et al., 2014; Nouws et al., 2023).

For the *Pseudomonas* sp. isolates, long-read data were re-processed on the Galaxy platform using two publicly available, Nanopore-optimised workflows. The Gene-Based Pathogen Identification workflow was reused for taxonomic consistency, while the Nanopore AMR Detection workflow was incorporated for dual consensus resistance profiling (Galaxy.eu, 2025). These workflows, adapted as needed, supported a final species-level characterisation that included refined taxonomic identification, de novo assembly, and the detection of virulence and resistance determinants.

### 2.7 Workflow outcome assessment

Workflow performance was evaluated by categorising each run into three discrete outcome states based on module execution status. A workflow was considered a complete success when all analytical modules (quality control, adapter trimming, read filtering, taxonomic classification, genome assembly, antimicrobial resistance, virulence factor, and plasmid detection) executed without premature termination. A partial success was defined as completing taxonomic assignment, with subsequent module failures in assembly, AMR, virulence, or plasmid detection. A complete failure occurred when upstream modules, such as quality control or read trimming, failed to complete.

### 2.8 Sequencing and assembly metrics

Read-level metrics (total reads, mean Q-score, and read N50) were computed for all sequencing runs passing basecalling quality filters. Assembly-level metrics (assembly N50 and contig counts) were compared only for datasets generated with extraction protocols that achieved partial (assembly-only) and complete workflow success, defined as uninterrupted execution through de novo assembly and downstream annotation modules.

### 2.9 Operational feasibility

Operational feasibility was evaluated for each extraction protocol across four domains: turnaround time (TAT), equipment portability, field suitability, and cost. TAT was defined as the duration of the extraction procedure. Hands-on and total elapsed TAT were recorded separately for each extraction and summarised by protocol.

Equipment portability was scored on a 1–10 scale, where higher values indicated compatibility with compact, potentially battery-operated devices. Scores considered the need for high-speed versus low-speed centrifugation, the possibility of using a simple heat block in place of a thermal shaker, and any requirement for specialised accessories such as magnetic racks. Field suitability reflected reagent storage conditions, tolerance to elevated ambient temperatures (30 °C), and procedural complexity, with higher ratings assigned to protocols that were cold chain sparing and operationally simple. Cost was assessed qualitatively on a per-sample basis from reagent and consumable prices (including Proteinase K where applicable) and reported as approximate per-sample cost in US dollars.

### 2.10 Statistical analysis

All statistical analyses were performed in R Studio v2025.9.2.418 (Posit team, 2025). Mean DNA yield and purity ratios (A260/A280) with standard deviations were calculated for each isolate and extraction protocol. Read-level metrics were compared across all extraction protocols using the Kruskal-Wallis test. In contrast, assembly contiguity metrics (assembly N50 and contig count) were compared only among protocols that produced successful assemblies for a given isolate set, using the Mann-Whitney U and Dunn-Holm pairwise testing for pairwise comparisons. Workflow completion rates (complete, partial, fail) by protocol were evaluated using Fisher’s exact test. The association between DNA purity and workflow success category was assessed with Spearman’s rank correlation coefficient (rho). Statistical significance was defined as p < 0.05.

## 3.0 RESULTS

### 3.1 DNA Yield and Purity Across Extraction Protocols

Across isolates, SwiftX DNA and SwiftX DNA + ProtK produced the highest DNA concentrations, typically ranging from 40 to 60 ng/µL for each sample. SwiftX ParaBact yielded lower concentrations, ranging from approximately 11 to 20 ng/µL, whereas NucleoSpin Microbial generated intermediate to high concentrations, with values up to about 100 ng/µL for *E. coli* TZ 3638 and CC-KAS-001-M1.

A260/280 purity ratios also varied by extraction protocol. SwiftX DNA and SwiftX DNA + ProtK showed ratios near 1.0–1.3 across isolates, SwiftX ParaBact produced ratios around 1.7–1.8, and NucleoSpin Microbial showed the highest and most uniform values, approximately 1.9–2.0 for all isolates. Detailed DNA yield and purity metrics for all 24 extracts are provided in Supplementary Table S1.

### 3.2 Sequencing performance

The extraction protocols had a marked impact on sequencing yield and read quality (Figure 2). For every isolate, the NucleoSpin Microbial kit generated the greatest total number of nanopore reads on a log scale (Figure 2a). SwiftX ParaBact produced intermediate read counts, and both SwiftX DNA-based approaches yielded the lowest output, indicating clear protocol-dependent differences in sequencing efficiency.

Read length and quality distributions further favoured the NucleoSpin Microbial approach. NucleoSpin Microbial showed the highest N50 read lengths (Figure 2b) and mean Q-scores (Figure 2c), SwiftX ParaBact produced moderately long reads with tightly clustered Q-scores, and SwiftX DNA and SwiftX DNA + ProtK were characterised by shorter N50 values and lower mean Q-scores, with Dunn-Holm pairwise testing confirming significant gains in read quality and smaller but detectable gains in N50 read length for NucleoSpin Microbial compared with the SwiftX protocols.

**Figure 2.**
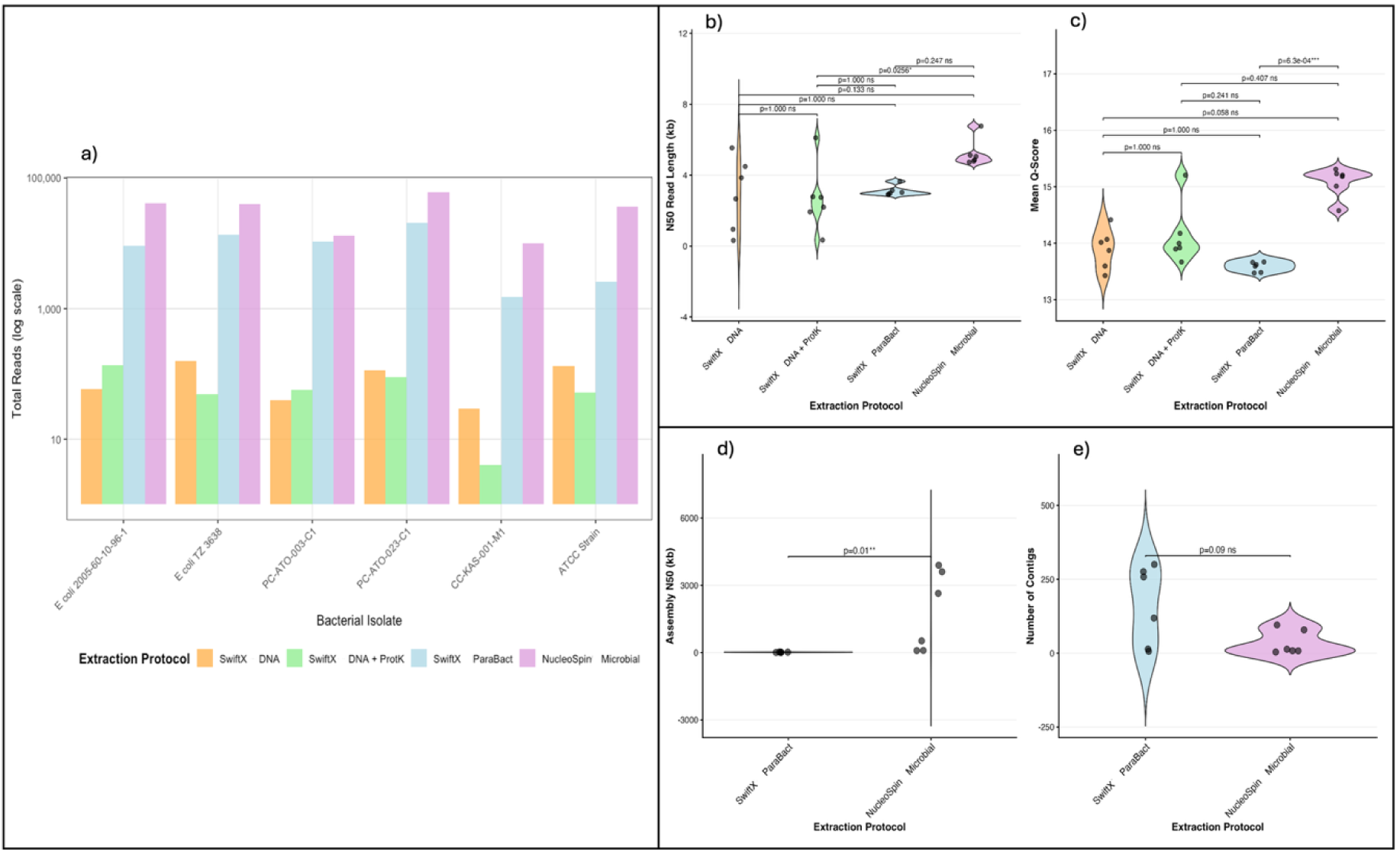
Sequencing and assembly performance metrics. (**a**) Total nanopore reads (log10 scale) per bacterial isolate for each DNA extraction protocol (SwiftX DNA, SwiftX DNA + ProtK, SwiftX ParaBact, NucleoSpin Microbial). NucleoSpin Microbial consistently yielded the highest read counts across all isolates. (**b, c**) Violin plots depicting read-level quality metrics: N50 read length (**b**) and mean Q-score (**c**) distributions across all four extraction protocols. Pairwise Dunn tests with Holm adjustment (p-values displayed above brackets) indicate statistically significant advantages for NucleoSpin Microbial in read quality. (**d, e**) Genome assembly contiguity metrics comparing the two protocols that successfully supported downstream assembly (SwiftX ParaBact and NucleoSpin Microbial). Panel ( **d**) shows Assembly N50 (kb), where NucleoSpin Microbial achieved significantly higher contiguity (p=0.01), and panel (**e**) shows the number of contigs per assembly. Individual points represent assemblies from independent isolates.

### 3.3 Species identification and taxonomic profiling

Using Kraken 2 for taxonomic profiling within the adapted Galaxy.eu workflow (Batut et al., 2025; Hiltemann et al., 2023), achieved perfect accuracy in identifying the predefined target organisms across all 24 samples from the four extraction protocols. The target-species read proportions for *E. coli* isolates ranged from 48.2% to 72.2%, while *Pseudomonas otitidis* and *Salmonella enterica* showed ranges of 79.3% to 100% and 45.4% to 66.4%, respectively. Taxonomic classification results confirming the identity of all six isolates are listed in Supplementary Table S2.

### 3.4 Bioinformatics workflow completion

Bioinformatics workflow completion varied substantially across extraction protocols. Complete end-to-end analysis was achieved in 100% (6/6) of isolates processed with the NucleoSpin Microbial and 83% (5/6) with SwiftX ParaBact protocols. In contrast, both the SwiftX DNA and SwiftX DNA + ProtK protocols yielded a 0% (0/6) completion rate. Details of the workflow completion rates by protocol and isolate are available in Supplementary Table S3.

### 3.5 Relationship between DNA purity and workflow success

Mean DNA purity (260/280 ratio) increased progressively from workflows that failed, through partially successful workflows, to those that achieved complete success (Figure 3). Failed workflows clustered at lower purity values (∼1.1–1.3), whereas complete workflows were consistently associated with higher purity ratios (∼1.7–2.0), with partial successes occupying an intermediate range, suggesting a threshold effect. Spearman’s rank correlation analysis confirmed a strong and statistically significant association between DNA purity and workflow outcome (Spearman’s ρ = 0.767, p < 0.0001), indicating that higher DNA purity strongly predicts successful completion of the sequencing and bioinformatics pipeline. Consistent with this trend, Kruskal–Wallis testing showed a significant increase in DNA purity across outcome categories from failed to fully successful workflows (p = 0.00059; Supplementary Figure S1). Protocol/kit-stratified scatterplots for this relationship are provided in Supplementary Figure S2.

**Figure 3.**
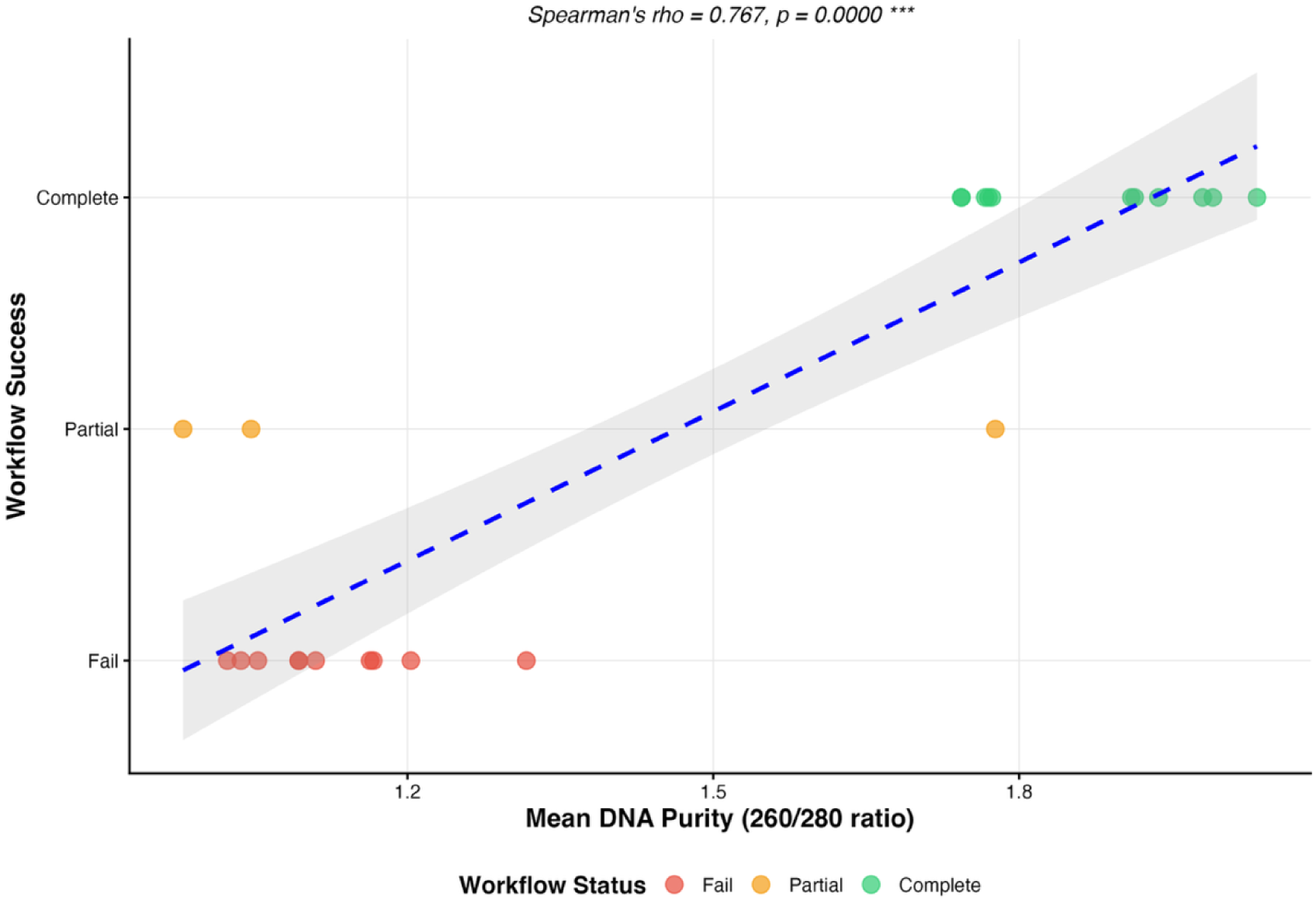
Correlation between DNA purity and downstream workflow success. Scatter plot showing the correlation between mean DNA purity, measured as the absorbance at 260/280 ratio, and overall workflow success. Each point represents an individual sample, coloured by workflow outcome (Fail, Partial, or Complete). The dashed blue line indicates the fitted trend, with the shaded area representing the 95% confidence interval. A strong positive correlation was observed between DNA purity and workflow success (Spearman’s ρ = 0.767, p < 0.0001), indicating that higher DNA purity is associated with an increased likelihood of successful completion of downstream sequencing and analysis workflows.

### 3.6 Failure mode characterisation

Failure modes within the bioinformatics workflow were not uniformly distributed across analysis stages (Supplementary Figure S3). In both SwiftX DNA and SwiftX DNA + ProtK protocols, all six isolates consistently failed at each major step, from initial quality control through plasmid detection, indicating a systemic upstream issue that prevented the species-specific pipelines from progressing. In contrast, SwiftX ParaBact extractions yielded no failures during quality control, trimming, filtering, taxonomic profiling, de novo assembly, polishing, and detection of virulence factors, with only isolated failures observed during AMR gene and plasmid detection. NucleoSpin Microbial extractions showed no failures at any stage, supporting the robustness of this protocol across the full analysis workflow.

### 3.7 Performance using species-specific bioinformatics pipelines (successful protocols only)

#### 3.7.1 Assembly contiguity

Across isolates, assemblies generated from NucleoSpin Microbial DNA showed higher N50 values than those from SwiftX ParaBact, with N50s clustering several kilobases higher than those from SwiftX ParaBact, and a significant difference between protocols (p = 0.01; Figure 2d). The number of contigs per assembly overlapped across protocols, with SwiftX ParaBact exhibiting a wider spread and NucleoSpin Microbial showing a slightly lower, more compact distribution (p = 0.09, not significant; Figure 2e).

#### 3.7.2 Virulence factor characterisation

Across all organism groups, the NucleoSpin Microbial protocol detected substantially higher numbers of both total and unique virulence factors than the SwiftX ParaBact protocol. Cumulatively, NucleoSpin Microbial accounted for 318 of 373 unique virulence factors identified (85.3%) and 365 of 446 total virulence factor detections (81.8%), whereas SwiftX ParaBact recovered only 55 unique virulence factors (14.7%) and 81 total detections (18.2%). The largest differences were observed for *Pseudomonas* sp. and *Salmonella* sp., for which NucleoSpin Microbial captured nearly the entire repertoire of detected virulence factors, while SwiftX ParaBact only captured a limited subset. At the isolate level, NucleoSpin Microbial yielded higher virulence factor counts (Supplementary Figure S4), reflected by a higher mean number of detections per isolate (60.8 ± 54.5) compared to SwiftX ParaBact (13.5 ± 12.7) (Supplementary Table S4). A comprehensive inventory of virulence factor genes detected by each protocol is provided in the Supplementary File S2.

#### 3.7.3 Pathogenicity Island detection

NucleoSpin Microbial showed broader and more consistent detection of pathogenicity islands. Strong connections were observed between NucleoSpin Microbial and SPI-1 through SPI-14, indicating successful recovery of most recognised SPIs, as well as the CS54 island (Supplementary Figure S5). The SwiftX ParaBact protocol, on the other hand, was only able to recover the SPI-14.

#### 3.7.4 Plasmid detection

The NucleoSpin Microbial and SwiftX ParaBact protocols both demonstrated high concordance, yielding identical plasmid profiles for five of the six bacterial isolates tested. The most complex plasmid profiles were observed in the *E. coli* isolates, with both extraction methods successfully recovering high-molecular-weight incompatibility (Inc) groups. Specifically, *E. coli* 2005-60-10-96-1 exhibited five distinct plasmid markers (IncFIA, IncFIB(AP001918), IncFIC, IncFII, and rep_cluster_2235), while *E. coli* TZ 3638 displayed the highest diversity with six detected targets, including IncQ1 and multiple replication clusters (rep_cluster_2244, rep_cluster_2350).

A notable variation was observed for isolate PC-ATO-003-C1. While the NucleoSpin Microbial protocol facilitated the detection of two distinct plasmid markers (rep_cluster_2244 and rep_cluster_2235), the SwiftX ParaBact method recovered only rep_cluster_2235. For the remaining isolates, results were consistent across both methods. The ATCC strain was confirmed to carry IncX1 and IncX3 plasmids, while no plasmid markers were detected in isolates PC-ATO-023-C1 or CC-KAS-001-M1 by either extraction protocol. Plasmid detection profiles for all six isolates across the two extraction protocols are detailed in Supplementary Figure S6.

#### 3.7.5 AMR gene detection

Across all isolates (Figure 4), the NucleoSpin Microbial protocol yielded a richer and more complex antimicrobial resistance (AMR) gene repertoire than SwiftX ParaBact, with numerous additional determinants detected for β-lactams (including several blaGES variants, blaOXA-10 and blaPOM-1), aminoglycosides (multiple aac and aph alleles), sulfonamides (sul1–3), trimethoprim (dfr genes), macrolides and tetracycline (tetA). Several multidrug-resistance patterns were only observed in assemblies derived from NucleoSpin Microbial DNA, whereas the corresponding SwiftX ParaBact assemblies frequently lacked these determinants or showed single-class resistance profiles. SwiftX ParaBact primarily recovered a narrower subset of AMR genes, most consistently sul, dfrA, aadA, and selected aph genes, and these were detected in fewer isolates.

**Figure 4.**
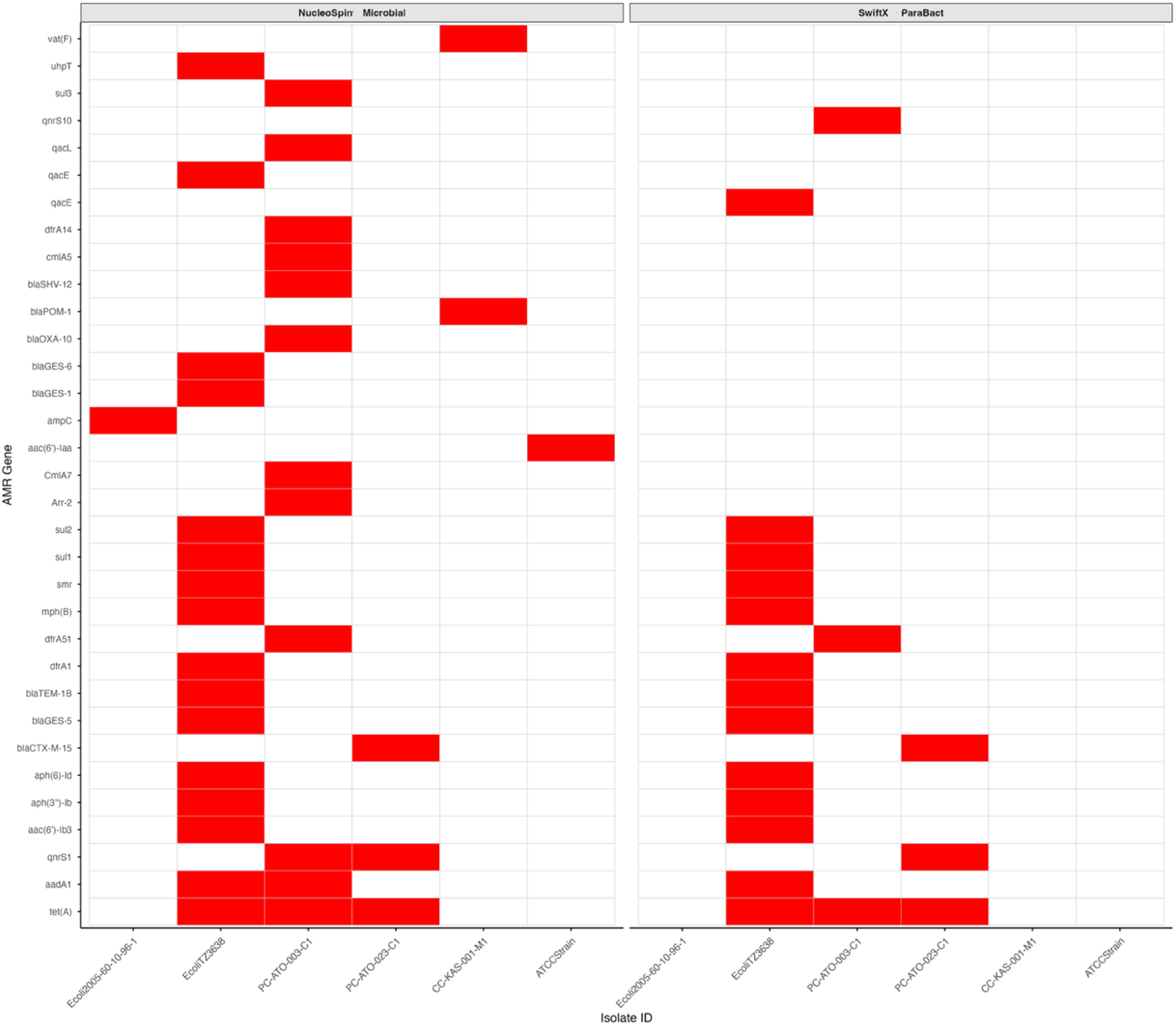
Antimicrobial resistance genes detected by each extraction protocol. Heatmap depicting presence (red) or absence (white) of AMR genes in bacterial isolates sequenced after DNA extraction with NucleoSpin Microbial (left panel) or SwiftX ParaBact (right panel); rows correspond to individual AMR genes and columns to isolates, enabling visual comparison of resistome breadth and revealing systematically higher AMR gene detection with NucleoSpin Microbial.

### 3.8 Operational performance and field-deployability assessment

The operational characteristics and field-deployability of the four DNA extraction protocols were evaluated across key criteria, including turnaround time, equipment portability, field suitability, cost, and technical complexity (Table 2). Turnaround times were shortest for SwiftX DNA (∼10 min hands-on, ∼15 min total), followed by SwiftX ParaBact (∼10 min hands-on, ∼20 min total) and SwiftX DNA + ProtK (∼15 min hands-on, ∼25 min total). NucleoSpin Microbial required the longest times (∼25 min hands-on, ∼35 min total). All protocols could be completed in less than 40 minutes from colony to purified DNA.

Equipment requirements translated into high equipment-portability scores for the SwiftX kits and a lower score for NucleoSpin Microbial. SwiftX DNA, SwiftX + ProtK, and SwiftX ParaBact were compatible with low-speed, portable mini-centrifuges and required only a simple heat block or thermal shaker and a magnetic rack, yielding portability scores of 9, 8, and 9 on a 10-point scale. In contrast, the NucleoSpin Microbial protocol required multiple 11,000× g centrifugation steps on a benchtop microcentrifuge but no magnetic rack, resulting in a portability score of 5.

Reagent storage requirements were least restrictive for SwiftX ParaBact (2–25 °C acceptable) and NucleoSpin Microbial (room temperature-stable kit, with Proteinase K stored at 4 °C or −20 °C after initial use). SwiftX DNA preferred 2–8 °C storage but tolerated up to 25 °C for ≤8 weeks, whereas SwiftX DNA + ProtK required −20 °C storage for Proteinase K plus chilled transport. At 30 °C, field stability was rated excellent for SwiftX DNA and SwiftX ParaBact, good for NucleoSpin Microbial, and moderate for SwiftX DNA + ProtK. Overall field suitability scores (1–10 scale) were highest for SwiftX ParaBact (9), followed by SwiftX DNA (8), NucleoSpin Microbial (7), and SwiftX DNA + ProtK (6).

Per-sample costs were low and comparable across protocols, ranging from about 3.30 USD for SwiftX DNA to 5.08 USD for NucleoSpin Microbial, with SwiftX + ProtK and SwiftX ParaBact falling in the intermediate range (approximately 4.80 and 4.30 USD per sample, respectively). All four protocols could be implemented with minimal training, with operators requiring less than 2 hours of hands-on familiarisation to perform the procedures independently.

**Table 2.**
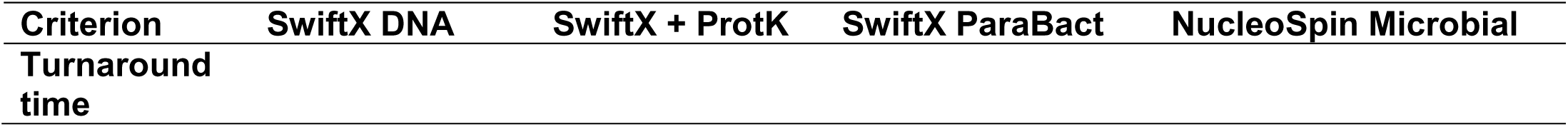
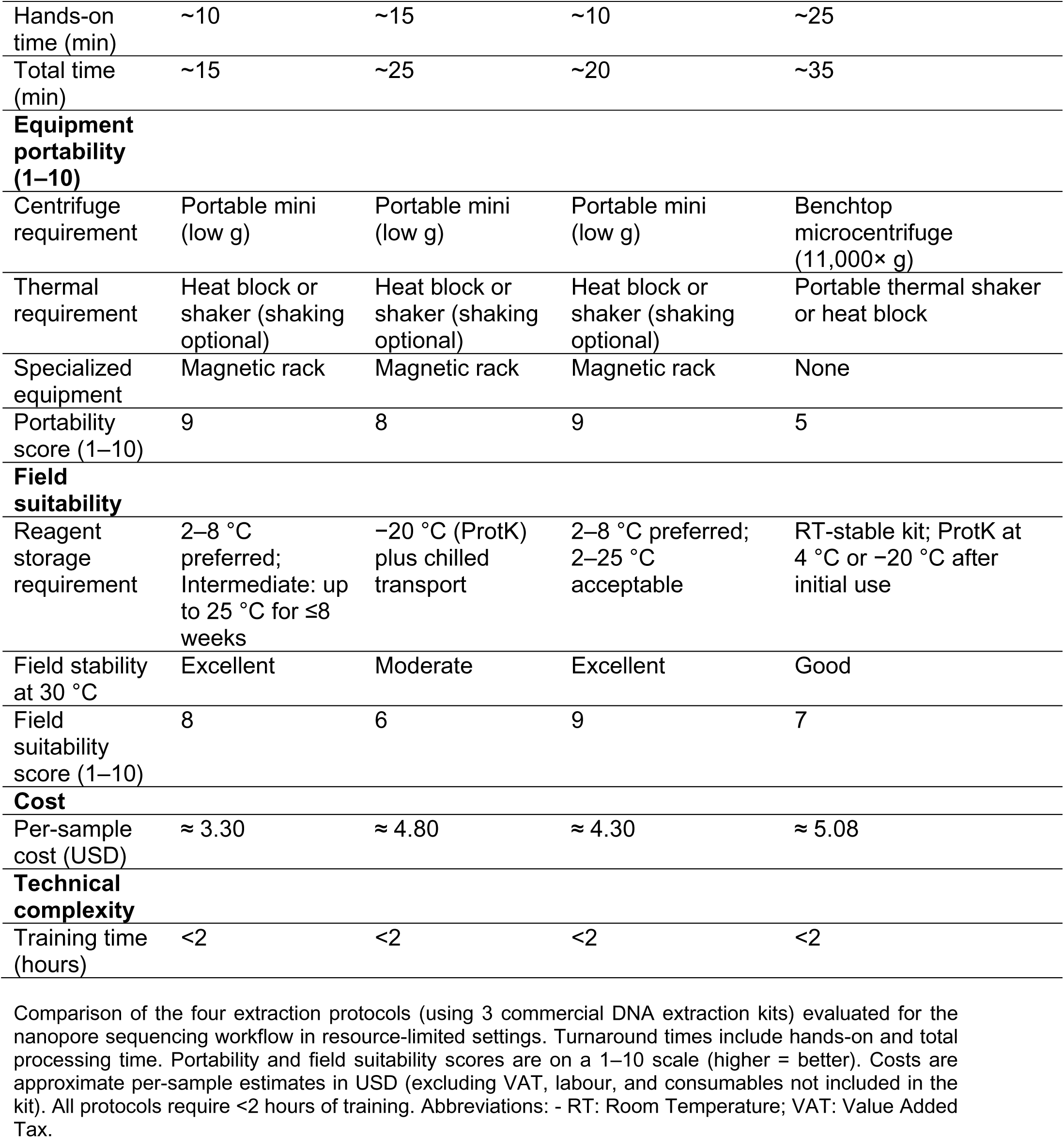
Operational performance and field-deployability assessment of DNA extraction protocols.

| Criterion | SwiftX DNA | SwiftX + ProtK | SwiftX ParaBact | NucleoSpin Microbial |
| --- | --- | --- | --- | --- |
| Turnaround time |  |  |  |  |
| Hands-on time (min) | ~10 | ~15 | ~10 | ~25 |
| Total time (min) | ~15 | ~25 | ~20 | ~35 |
| **Equipment portability (1–10)** |  |  |  |  |
| Centrifuge requirement | Portable mini (low g) | Portable mini (low g) | Portable mini (low g) | Benchtop microcentrifuge (11,000× g) |
| Thermal requirement | Heat block or shaker (shaking optional) | Heat block or shaker (shaking optional) | Heat block or shaker (shaking optional) | Portable thermal shaker or heat block |
| Specialized equipment | Magnetic rack | Magnetic rack | Magnetic rack | None |
| Portability score (1–10) | 9 | 8 | 9 | 5 |
| **Field suitability** |  |  |  |  |
| Reagent storage requirement | 2–8 °C preferred; Intermediate: up to 25 °C for ≤8 weeks | –20 °C (ProtK) plus chilled transport | 2–8 °C preferred; 2–25 °C acceptable | RT-stable kit; ProtK at 4 °C or –20 °C after initial use |
| Field stability at 30 °C | Excellent | Moderate | Excellent | Good |
| Field suitability score (1–10) | 8 | 6 | 9 | 7 |
| **Cost** |  |  |  |  |
| Per-sample cost (USD) | ≈ 3.30 | ≈ 4.80 | ≈ 4.30 | ≈ 5.08 |
| **Technical complexity** |  |  |  |  |
| Training time (hours) | <2 | <2 | <2 | <2 |

Comparison of the four extraction protocols (using 3 commercial DNA extraction kits) evaluated for the nanopore sequencing workflow in resource-limited settings. Turnaround times include hands-on and total processing time. Portability and field suitability scores are on a 1–10 scale (higher = better). Costs are approximate per-sample estimates in USD (excluding VAT, labour, and consumables not included in the kit). All protocols require <2 hours of training. Abbreviations: - RT: Room Temperature; VAT: Value Added Tax.

## 4.0 DISCUSSION

### 4.1 DNA Purity as a critical determinant of nanopore sequencing success

This study presents a comprehensive comparison of portable DNA extraction protocols for Oxford Nanopore sequencing, explicitly linking extraction chemistry to the successful completion of downstream bioinformatics workflows. While previous studies have independently evaluated extraction yield or sequencing metrics (Peker et al., 2021; Purushothaman et al., 2024; Thornval et al., 2025), few have demonstrated how extraction quality directly determines the feasibility of completing clinically relevant genomic pipelines, particularly in low- and middle-income countries (LMIC) contexts.

We observed a strong and statistically significant association between DNA purity and successful completion of downstream bioinformatics workflows. Samples with purity values in the optimal range (∼1.8–2.0) consistently completed all analytical stages, whereas samples with lower ratios (≤1.3) failed early, typically during quality control or shortly after taxonomic assignment. This is consistent with Oxford Nanopore Technologies’ guidance, which emphasises high spectrophotometric purity for stable sequencing performance (Oxford Nanopore Technologies, 2025; Koetsier and Cantor, 2019).

Importantly, our findings extend this guidance by demonstrating that DNA extracts deemed acceptable by conventional spectrophotometric or qPCR-based metrics, as reported by Antane and colleagues, can still compromise nanopore read quality and hinder the completion of clinically relevant, species-specific pipelines (Antane et al., 2024). The strong correlation observed between DNA purity and workflow success (Spearman’s ρ = 0.767, p < 0.0001) indicates that purity metrics provide a practical predictor of pipeline robustness and a useful pre-sequencing screening tool. These results are particularly relevant for resource-limited settings, where early identification of suboptimal extracts could reduce failed runs, conserve sequencing resources, and improve the reliability of field-deployable genomic diagnostics (Pronyk et al., 2023; Wasswa et al., 2022).

### 4.2 Protocol-dependent differences in sequencing output and assembly quality

Sequencing performance varied significantly between extraction protocols, consistent with previous comparisons of long-read workflows and DNA isolation methods (Bari et al., 2025; Peker et al., 2021; Schurig et al., 2024). NucleoSpin Microbial, which was the reference gold standard in this study, consistently achieved the highest read yields, longest N50 values, and highest mean Q-scores across all isolates, resulting in superior assemblies. SwiftX DNA and SwiftX DNA + ProtK, designed as rapid “direct extraction” workflows with minimal purification, produced very low sequencing outputs and frequent workflow failures, consistent with evidence that limited clean-up impairs downstream amplification and sequencing even when lysis is efficient (Jones et al., 2021; Sidstedt et al., 2020). SwiftX ParaBact, despite also using paramagnetic bead-based reverse purification, performed comparably to NucleoSpin Microbial with robust read quality and assembly metrics. This suggests that the SwiftX ParaBact kit mitigates the limitations of direct extraction approaches. NucleoSpin Microbial’s combination of mechanical (glass bead beating), enzymatic lysis, and silica-column purification provided marginal additional benefits, like in other studies by Purushothaman et al. (2024), for the most comprehensive recovery of virulence factors, AMR genes and plasmid regions, especially in *Pseudomonas* and *Salmonella* isolates (Purushothaman et al., 2024).

### 4.3 Practical implications for virulence factor and pathogenicity island detection

The marked differences in virulence factor and pathogenicity island recovery observed between extraction protocols (Supplementary Table S4 & Figure S5) demonstrate that DNA quality has direct consequences for how completely key pathogenicity determinants are represented in genome assemblies (Schmidt & Hensel, 2004). NucleoSpin Microbial provided the most exhaustive recovery of virulence genes and *Salmonella* pathogenicity islands, supporting a detailed assessment of invasion, survival, and transmission potential, whereas SwiftX ParaBact yielded a more limited but still informative subset of these loci, sufficient for broad inference of pathogenic potential in many surveillance and diagnostic contexts.

### 4.4 Plasmid detection and antimicrobial resistance gene diversity

Plasmid and AMR profiling showed that both extraction workflows recovered the clinically important plasmid backbones that commonly mobilise extended-spectrum beta-lactamases (ESBLs), carbapenemases and other priority resistance determinants in Enterobacterales, including IncFIA, IncFIB(AP001918), IncFII, IncFIC, IncQ1, IncX1, IncX3 and several rep clusters (Carattoli et al., 2014; Carattoli & Hasman, 2020). This incompatibility spectrum mirrors global data indicating that IncF and IncX plasmids dominate the dissemination of CTX-M ESBLs and NDM-type carbapenemases across clinical and food-chain settings (Mathers et al., 2015; Rozwandowicz et al., 2018).

A critical discrepancy, however, emerged in the recovery of the associated antimicrobial resistance (AMR) cargo genes. The NucleoSpin Microbial revealed extensive multidrug resistance profiles, including blaGES, blaPOM-1, multiple aac/aph alleles, sul1–3, dfr and tetA, whereas many of these determinants were missed with SwiftX ParaBact extraction. This “gene drop-out” phenomenon of detecting the plasmid backbone but missing the resistance gene underpins how extraction choice profoundly shapes the apparent resistome (Chow et al., 2001; Murray et al., 2022; Papamentzelopoulou et al., 2025). Likely, the mechanical bead-beating and enzymatic digestion employed by the NucleoSpin Microbial kit are necessary to fully lyse cells containing high-copy resistance plasmids or to preserve the integrity of specific plasmid regions during purification. Our findings align with previous studies, highlighting the critical role of combining mechanical bead beating with enzymatic lysis to achieve efficient DNA recovery (Kruasuwan et al., 2024). Nonetheless, SwiftX ParaBact reliably captured the core IncF/IncX backbones and key resistance genes sufficient to classify isolates as multidrug-resistant, supporting its use for frontline surveillance, with referral of flagged isolates to higher-resolution workflows.

### 4.5 Operational feasibility and field deployability

Despite the superior sequencing and diagnostic performance of NucleoSpin Microbial, the SwiftX protocols offer notable practical advantages for field deployment. All three SwiftX variants run on low-speed mini-centrifuges, simple heat blocks, and magnetic racks, with total extraction times of about 10–20 minutes versus roughly 35 minutes for NucleoSpin Microbial. SwiftX ParaBact reagents are stable at 2–25 °C, whereas components of NucleoSpin Microbial (for example, Proteinase K) require refrigerated or frozen storage, a major constraint where cold-chain infrastructure is unreliable (Dauner et al., 2015).

These findings highlight a practical trade-off: magnetic bead–based methods such as SwiftX ParaBact maximise portability and simplicity, whereas silica-column extraction with NucleoSpin Microbial maximises diagnostic completeness and bioinformatics robustness. A realistic approach for resource-limited surveillance systems is a two-tier model, using SwiftX for rapid field screening and reserving NucleoSpin for confirmatory work at reference laboratories, while future optimisation of reverse-purification chemistries may narrow this performance gap.

### 4.6 Strengths and limitations

The strengths of this study include the use of a phylogenetically diverse bacterial panel, the direct link between extraction performance and complete bioinformatics workflow outcomes (rather than isolated metrics such as read count alone), and the practical focus on operational feasibility in resource-limited contexts. The use of NucleoSpin Microbial as a reference laboratory gold standard for analytical completeness and accuracy provides a standardised benchmark against which rapid field-deployable methods can be evaluated (Besset-Manzoni et al., 2019; Laganenka et al., 2019; Pheiffer et al., 2022; Prieto-Fernández et al., 2024). The multiplexing of all 24 extracts on a single sequencing run minimised batch effects, ensuring that observed differences were attributable to the extraction protocols. Furthermore, the employment of validated, species-specific bioinformatics pipelines (STEC v1.2, Salmonella v1.0, Nanopore AMR workflows) strengthens the clinical relevance of the findings (Abueg et al., 2024; Jaudou et al., 2022; Timilsina et al., 2025).

Limitations include the modest sample size (six isolates) and the focus on only three target species. Expansion to additional pathogenic bacteria (e.g., *Mycobacterium tuberculosis*, *Klebsiella pneumoniae*, *Acinetobacter baumannii*) would strengthen the generalizability of findings. Additionally, the study was restricted to three commercial extraction kits, specifically selected for their portability, accessibility, and minimal equipment requirements, rather than providing an exhaustive market survey. Conventional magnetic bead–based extraction methods were not included. Although the magnetic bead approach for nucleic acid extraction is simple to operate, several critical operational details must be optimised to ensure consistent yield and purity, including the number of magnetic beads, the number of wash cycles, dissociation/elution conditions, and pipetting/mixing dynamics during binding and washing steps (He et al., 2017; Lee et al., 2023). This makes them less practical for field deployment.

Finally, the evaluation of reagent field stability was qualitative, and the study utilised R10.4.1 flow cells; future work should incorporate quantitative environmental stress testing and comparison with newer pore generations to further contextualise these results.

## 5.0 CONCLUSION

This comparative evaluation study shows that DNA extraction is a key driver of nanopore diagnostic performance, with DNA purity strongly predicting full workflow completion (ρ = 0.767, p < 0.0001). NucleoSpin Microbial delivered 100% workflow success and the most complete virulence, plasmid, and AMR profiles, making it the preferred option for centralised laboratories that require maximal diagnostic accuracy. SwiftX ParaBact provided a useful middle ground, offering high purity, 83% workflow success, and recovery of core virulence and resistance determinants suitable for many surveillance applications. Although SwiftX DNA-based protocols failed at assembly and detailed AMR profiling, they consistently supported correct taxonomic profiling and therefore retain value for rapid species-level screening. Programmes planning nanopore-based AMR surveillance in resource-limited settings should therefore prioritise optimisation and selection of extraction workflows, with future work focusing on improved or hybrid bead/column chemistries that combine portability with high analytical performance.

## 6.0 AUTHOR CONTRIBUTIONS

**LK-T**: Conceptualization, Data curation, Investigation, Formal analysis, Methodology, Visualisation, Writing – original draft, Writing – review & editing. **SKA & AKA**: Data curation, Formal analysis, Investigation, Methodology, Writing – review & editing. **DSB & WNSA**: Data curation, Investigation, Writing – review & editing. **VNBF**: Data curation, Formal analysis, Writing – review & editing. **RNF**: Project administration, Writing – review & editing. **FA-Y & RMK:** Conceptualization, Methodology, Supervision, Writing – review & editing. **UT**, **AAEW, YAA, & ROP**: Conceptualization, Funding acquisition, Resources, Methodology, Supervision, Writing – review & editing. **MF**: Conceptualization, Data curation, Investigation, Formal analysis, Validation, Methodology, Funding acquisition, Resources, Supervision, Writing – review & editing.

## 9.0 ETHICAL APPROVAL

Ethical approval (Ref: CHRPE/AP/446/24) was obtained from the Committee on Human Research Publications and Ethics of the Kwame Nkrumah University of Science and Technology.

## ACKNOWLEDGEMENTS

We would like to appreciate the German Federal Institute for Risk Assessment (BfR) for securing some of the isolates used in this study from the European Reference Laboratory for Antimicrobial Resistance (EURL-AR) in DTU. We also extend our gratitude to the One Health Bacteriology group and the logistics department at the Kumasi Centre for Collaborative Research in Tropical Medicine (KCCR). We further acknowledge the African One Health Network for Disease Prevention (ADAPT consortium) for their contributions to this study.

## REFERENCES

Abueg, L. A. L., Afgan, E., Allart, O., Awan, A. H., Bacon, W. A., Baker, D., Bassetti, M., Batut, B., Bernt, M., Blankenberg, D., Bombarely, A., Bretaudeau, A., Bromhead, C. J., Burke, M. L., Capon, P. K., Čech, M., Chavero-Díez, M., Chilton, J. M., Collins, T. J., … Zoabi, R. (2024). The Galaxy platform for accessible, reproducible, and collaborative data analyses: 2024 update. Nucleic Acids Research, 52(W1), W83–W94. 10.1093/NAR/GKAE410

Antane, V., Sarybayev, Y., Osserbay, A., Shatmanov, K., & Baltakhozhayev, T. (2024). Spectrophotometric method for determining the quantity and quality of DNA in animal breeding. Scientific Horizons, 27(2), 31–42. 10.48077/SCIHOR2.2024.31

Aruhomukama, D. (2022). Antimicrobial resistance data, frugal sequencing, and low-income countries in Africa. The Lancet Infectious Diseases, 22(7), 933–934. 10.1016/S1473-3099(22)00312-7

Bankevich, A., Nurk, S., Antipov, D., Gurevich, A. A., Dvorkin, M., Kulikov, A. S., Lesin, V. M., Nikolenko, S. I., Pham, S., Prjibelski, A. D., Pyshkin, A. V., Sirotkin, A. V., Vyahhi, N., Tesler, G., Alekseyev, M. A., & Pevzner, P. A. (2012). SPAdes: A New Genome Assembly Algorithm and Its Applications to Single-Cell Sequencing. Https://Home.Liebertpub.Com/Cmb, 19(5), 455–477. 10.1089/CMB.2012.0021

Bari, A. K., Xavier, B. B., Severs, T., Sinha, B., & Rossen, J. W. A. (2025). Comparison of three commercial DNA extraction kits and assemblers for AMR determinant detection in Pseudomonas aeruginosa and Enterobacter cloacae using long-read sequencing. Journal of Microbiological Methods, 239, 107317. 10.1016/J.MIMET.2025.107317

Batut, B., Nasr, E., & Zierep, P. (2025). Pathogen detection from (direct Nanopore) sequencing data using Galaxy-Foodborne Edition.

Besset-Manzoni, Y., Joly, P., Brutel, A., Gerin, F., Soudière, O., Langin, T., & Prigent-Combaret, C. (2019). Does in vitro selection of biocontrol agents guarantee success in planta? A study case of wheat protection against Fusarium seedling blight by soil bacteria. PLOS ONE, 14(12), e0225655. 10.1371/JOURNAL.PONE.0225655

Bogaerts, B., Nouws, S., Verhaegen, B., Denayer, S., Van Braekel, J., Winand, R., Fu, Q., Crombé, F., Piérard, D., Marchal, K., C Roosens, N. H., J De Keersmaecker, S. C., & Vanneste, K. (2021). Validation strategy of a bioinformatics whole genome sequencing workflow for Shiga toxin-producing Escherichia coli using a reference collection extensively characterized with conventional methods DATA SUMMARY. Microbial Genomics, 7, 531. 10.1099/mgen.0.000531

Bogaerts, B., Van Braekel, J., Van Uffelen, A., D’aes, J., Godfroid, M., Delcourt, T., Kelchtermans, M., Milis, K., Goeders, N., De Keersmaecker, S. C. J., Roosens, N. H. C., Winand, R., & Vanneste, K. (2025). Galaxy @Sciensano: a comprehensive bioinformatics portal for genomics-based microbial typing, characterization, and outbreak detection. BMC Genomics, 26(1), 20. 10.1186/S12864-024-11182-5

Bortolaia, V., Kaas, R. S., Ruppe, E., Roberts, M. C., Schwarz, S., Cattoir, V., Philippon, A., Allesoe, R. L., Rebelo, A. R., Florensa, A. F., Fagelhauer, L., Chakraborty, T., Neumann, B., Werner, G., Bender, J. K., Stingl, K., Nguyen, M., Coppens, J., Xavier, B. B., … Aarestrup, F. M. (2020). ResFinder 4.0 for predictions of phenotypes from genotypes. Journal of Antimicrobial Chemotherapy, 75(12), 3491–3500. 10.1093/JAC/DKAA345

Byrne, R. L., Cocker, D., Alyayyoussi, G., Mphasa, M., Charles, M., Mandula, T., Williams, C. T., Rigby, J., Hearn, J., Feasey, N., Adams, E. R., & Edwards, T. (2022). A novel, magnetic bead-based extraction method for the isolation of antimicrobial resistance genes with a case study in river water in Malawi. Journal of Applied Microbiology, 133(5), 3191. 10.1111/JAM.15755

Carattoli, A., & Hasman, H. (2020). PlasmidFinder and In Silico pMLST: Identification and Typing of Plasmid Replicons in Whole-Genome Sequencing (WGS). Methods in Molecular Biology, 2075, 285–294. 10.1007/978-1-4939-9877-7_20

Carattoli, A., Zankari, E., Garciá-Fernández, A., Larsen, M. V., Lund, O., Villa, L., Aarestrup, F. M., & Hasman, H. (2014). In Silico Detection and Typing of Plasmids using PlasmidFinder and Plasmid Multilocus Sequence Typing. Antimicrobial Agents and Chemotherapy, 58(7), 3895. 10.1128/AAC.02412-14

Chen, L., Yang, J., Yu, J., Yao, Z., Sun, L., Shen, Y., & Jin, Q. (2004). VFDB: a reference database for bacterial virulence factors. Nucleic Acids Research, 33(Database Issue), D325. 10.1093/NAR/GKI008

Chen, S. (2023). Ultrafast one-pass FASTQ data preprocessing, quality control, and deduplication using fastp. IMeta, 2(2), e107. 10.1002/IMT2.107;JOURNAL:JOURNAL:2770596X

Chen, S., Zhou, Y., Chen, Y., & Gu, J. (2018). fastp *: an ultra-fast all-in-one FASTQ preprocessor*. 10.1101/274100

Chin, N. A., Salihah, N. T., Shivanand, P., & Ahmed, M. U. (2021). Recent trends and developments of PCR-based methods for the detection of food-borne Salmonella bacteria and Norovirus. Journal of Food Science and Technology, 59(12), 4570. 10.1007/S13197-021-05280-5

Chow, J. W., Kak, V., You, I., Kao, S. J., Petrin, J., Clewell, D. B., Lerner, S. A., Miller, G. H., & Shaw, K. J. (2001). Aminoglycoside Resistance Genes aph(2”)-Ib and aac(6′)-Im Detected Together in Strains of both Escherichia coli and Enterococcus faecium. Antimicrobial Agents and Chemotherapy, 45(10), 2691. 10.1128/AAC.45.10.2691-2694.2001

Dauner, A. L., Gilliland, T. C., Mitra, I., Pal, S., Morrison, A. C., Hontz, R. D., & Wu, S. J. L. (2015). Evaluation of Nucleic Acid Stabilization Products for Ambient Temperature Shipping and Storage of Viral RNA and Antibody in a Dried Whole Blood Format. The American Journal of Tropical Medicine and Hygiene, 93(1), 46–53. 10.4269/AJTMH.15-0110

De Coster, W., D’Hert, S., Schultz, D. T., Cruts, M., & Van Broeckhoven, C. (2018). NanoPack: visualizing and processing long-read sequencing data. Bioinformatics, 34(15), 2666–2669. 10.1093/BIOINFORMATICS/BTY149

Donkor, E. S., Odoom, A., Osman, A. H., Darkwah, S., & Kotey, F. C. N. (2024). A Systematic Review on Antimicrobial Resistance in Ghana from a One Health Perspective. Antibiotics, 13(7), 662. 10.3390/ANTIBIOTICS13070662/S1

Elbehiry, A., Marzouk, E., & Abalkhail, A. (2025). Antimicrobial resistance at a turning point: microbial drivers, one health, and global futures. Frontiers in Microbiology, 16, 1698809. 10.3389/FMICB.2025.1698809

Ewels, P., Magnusson, M., Lundin, S., & Käller, M. (2016). MultiQC: summarize analysis results for multiple tools and samples in a single report. Bioinformatics, 32(19), 3047–3048. 10.1093/BIOINFORMATICS/BTW354

Feldgarden, M., Brover, V., Haft, D. H., Prasad, A. B., Slotta, D. J., Tolstoy, I., Tyson, G. H., Zhao, S., Hsu, C. H., McDermott, P. F., Tadesse, D. A., Morales, C., Simmons, M., Tillman, G., Wasilenko, J., Folster, J. P., & Klimke, W. (2019). Validating the AMRFINder tool and resistance gene database by using antimicrobial resistance genotype-phenotype correlations in a collection of isolates. Antimicrobial Agents and Chemotherapy, 63(11). 10.1128/AAC.00483-19

*Galaxy*. (n.d.). Retrieved December 1, 2025, from https://usegalaxy.eu/workflows/list

*Galaxy | Galaxy External*. (n.d.). Retrieved November 30, 2025, from https://galaxy.sciensano.be/datasets/21d90a595e8c34b9/preview

*GitHub - nanoporetech/medaka: Sequence correction provided by ONT Research*. (n.d.). Retrieved November 30, 2025, from https://github.com/nanoporetech/medaka

Gurevich, A., Saveliev, V., Vyahhi, N., & Tesler, G. (2013). QUAST: quality assessment tool for genome assemblies. Bioinformatics, 29(8), 1072–1075. 10.1093/BIOINFORMATICS/BTT086

He, H., Li, R., Chen, Y., Pan, P., Tong, W., Dong, X., Chen, Y., & Yu, D. (2017). Integrated DNA and RNA extraction using magnetic beads from viral pathogens causing acute respiratory infections. Scientific Reports 2017 7:1, 7(1), 45199-. 10.1038/srep45199

Hiltemann, S., Rasche, H., Gladman, S., Hotz, H. R., Larivière, D., Blankenberg, D., Jagtap, P. D., Wollmann, T., Bretaudeau, A., Goué, N., Griffin, T. J., Royaux, C., Bras, Y. Le, Mehta, S., Syme, A., Coppens, F., Droesbeke, B., Soranzo, N., Bacon, W., … Batut, B. (2023). Galaxy Training: A powerful framework for teaching! PLOS Computational Biology, 19(1), e1010752. 10.1371/JOURNAL.PCBI.1010752

Hoenen, T., Groseth, A., Rosenke, K., Fischer, R. J., Hoenen, A., Judson, S. D., Martellaro, C., Falzarano, D., Marzi, A., Squires, R. B., Wollenberg, K. R., De Wit, E., Prescott, J., Safronetz, D., van Doremalen, N., Bushmaker, T., Feldmann, F., McNally, K., Bolay, F. K., … Feldmann, H. (2016). Nanopore sequencing as a rapidly deployable Ebola outbreak tool. Emerging Infectious Diseases, 22(2), 331–334. 10.3201/eid2202.151796

Inouye, M., Dashnow, H., Raven, L. A., Schultz, M. B., Pope, B. J., Tomita, T., Zobel, J., & Holt, K. E. (2014). SRST2: Rapid genomic surveillance for public health and hospital microbiology labs. Genome Medicine 2014 6:11, 6(11), 90-. 10.1186/S13073-014-0090-6

*Input DNA/RNA QC | Oxford Nanopore Technologies*. (n.d.). Retrieved December 18, 2025, from https://nanoporetech.com/document/input-dna-rna-qc

Jaudou, S., Deneke, C., Tran, M. L., Schuh, E., Goehler, A., Vorimore, F., Malorny, B., Fach, P., Grützke, J., & Delannoy, S. (2022). A step forward for Shiga toxin-producing Escherichia coli identification and characterization in raw milk using long-read metagenomics. Microbial Genomics, 8(11), mgen000911. 10.1099/MGEN.0.000911

Jaureguy, F., Landraud, L., Passet, V., Diancourt, L., Frapy, E., Guigon, G., Carbonnelle, E., Lortholary, O., Clermont, O., Denamur, E., Picard, B., Nassif, X., & Brisse, S. (2008). Phylogenetic and genomic diversity of human bacteremic Escherichia coli strains. BMC Genomics 2008 9:1, 9(1), 560-. 10.1186/1471-2164-9-560

Jin, Y., Li, Y., Huang, S., Hong, C., Feng, X., Cai, H., Xia, Y., Li, S., Zhang, L., Lou, Y., & Guan, W. (2024). Whole-Genome Sequencing Analysis of Antimicrobial Resistance, Virulence Factors, and Genetic Diversity of Salmonella from Wenzhou, China. Microorganisms, 12(11), 2166. 10.3390/MICROORGANISMS12112166/S1

Joensen, K. G., Scheutz, F., Lund, O., Hasman, H., Kaas, R. S., Nielsen, E. M., & Aarestrup, F. M. (2014). Real-time whole-genome sequencing for routine typing, surveillance, and outbreak detection of verotoxigenic Escherichia coli. Journal of Clinical Microbiology, 52(5), 1501–1510. 10.1128/JCM.03617-13;JOURNAL:JOURNAL:JCM;WGROUP:STRING:PUBLICATION

Joensen, K. G., Tetzschner, A. M. M., Iguchi, A., Aarestrup, F. M., & Scheutz, F. (2015). Rapid and easy in silico serotyping of Escherichia coli isolates by use of whole-genome sequencing data. Journal of Clinical Microbiology, 53(8), 2410–2426. 10.1128/JCM.00008-15;JOURNAL:JOURNAL:JCM;ISSUE:ISSUE:DOI

Jones, A., Torkel, C., Stanley, D., Nasim, J., Borevitz, J., & Schwessinger, B. (2021). High-molecular weight DNA extraction, clean-up and size selection for long-read sequencing. PLoS ONE, 16(7), e0253830. 10.1371/JOURNAL.PONE.0253830

Katz, L. S., Griswold, T., Lindsey, R. L., Lauer, A. C., Im, M. S., Williams, G., Halpin, J. L., Gómez, G. A., Kucerova, Z., Morrison, S., Page, A., Den Bakker, H. C., & Carleton, H. A. (2025). Kalamari: a representative set of genomes of public health concern. Microbiology Resource Announcements, 14(2). 10.1128/mra.00963-24

Koetsier, G., Cantor, E., & Biolabs, E. (n.d.). *A Practical Guide to Analyzing Nucleic Acid Concentration and Purity with Microvolume Spectrophotometers*.

Kolmogorov, M., Yuan, J., Lin, Y., & Pevzner, P. A. (2019). Assembly of long, error-prone reads using repeat graphs. Nature Biotechnology 2019 37:5, 37(5), 540–546. 10.1038/s41587-019-0072-8

Kombade, S., Kaur, N., Kombade, S., & Kaur, N. (2021). Pathogenicity Island in *Salmonella Spp**. -* A Global Challenge. 10.5772/INTECHOPEN.96443

Kruasuwan, W., Sawatwong, P., Jenjaroenpun, P., Wankaew, N., Arigul, T., Yongkiettrakul, S., Lunha, K., Sudjai, A., Siludjai, D., Skaggs, B., & Wongsurawat, T. (2024). Comparative evaluation of commercial DNA isolation approaches for nanopore-only bacterial genome assembly and plasmid recovery. Scientific Reports 2024 14:1, 14(1), 27672-. 10.1038/s41598-024-78066-2

Kuupiel, D., Bawontuo, V., & Mashamba-Thompson, T. P. (2017). Improving the Accessibility and Efficiency of Point-of-Care Diagnostics Services in Low- and Middle-Income Countries: Lean and Agile Supply Chain Management. Diagnostics 2017, Vol. 7, Page 58, 7(4), 58. 10.3390/DIAGNOSTICS7040058

Laganenka, L., Sander, T., Lagonenko, A., Chen, Y., Link, H., & Sourjik, V. (2019). Quorum sensing and metabolic state of the host control lysogeny-lysis switch of bacteriophage T1. MBio, 10(5). 10.1128/MBIO.0188419

Lamas, A., Garrido-Maestu, A., Prieto, A., Cepeda, A., & Franco, C. M. (2023). Whole genome sequencing in the palm of your hand: how to implement a MinION Galaxy-based workflow in a food safety laboratory for rapid Salmonella spp. serotyping, virulence, and antimicrobial resistance gene identification. Frontiers in Microbiology, 14, 1254692. 10.3389/FMICB.2023.1254692/FULL

Laxminarayan, R. (2022). The overlooked pandemic of antimicrobial resistance. The Lancet, 399(10325), 606–607. 10.1016/S0140-6736(22)00087-3

Lee, S. M., Balakrishnan, H. K., Doeven, E. H., Yuan, D., & Guijt, R. M. (2023). Chemical Trends in Sample Preparation for Nucleic Acid Amplification Testing (NAAT): A Review. Biosensors, 13(11), 980. 10.3390/BIOS13110980

Lerminiaux, N., Fakharuddin, K., Adam, H. J., Bharat, A., Golding, G. R., Martin, I., Mulvey, M., & Mataseje, L. (2025). Rapid identification of microbial pathogens and antimicrobial resistance from bloodstream infections using long-read sequencing. BioRxiv, 2025.12.12.694010. 10.64898/2025.12.12.694010

Li, H. (2021). New strategies to improve minimap2 alignment accuracy. Bioinformatics, 37(23), 4572–4574. 10.1093/BIOINFORMATICS/BTAB705

Li, H., Handsaker, B., Wysoker, A., Fennell, T., Ruan, J., Homer, N., Marth, G., Abecasis, G., & Durbin, R. (2009). The Sequence Alignment/Map format and SAMtools. Bioinformatics, 25(16), 2078–2079. 10.1093/BIOINFORMATICS/BTP352

Liu, L., Li, Y., Li, S., Hu, N., He, Y., Pong, R., Lin, D., Lu, L., & Law, M. (2012). Comparison of next-generation sequencing systems. Journal of Biomedicine & Biotechnology, 2012, 251364. 10.1155/2012/251364

Low, A. J., Koziol, A. G., Manninger, P. A., Blais, B., & Carrillo, C. D. (2019). ConFindr: Rapid detection of intraspecies and cross-species contamination in bacterial whole-genome sequence data. PeerJ, 2019(5), e6995. 10.7717/PEERJ.6995/SUPP-2

Mathers, A. J., Peirano, G., & Pitout, J. D. D. (2015). The Role of Epidemic Resistance Plasmids and International High-Risk Clones in the Spread of Multidrug-Resistant Enterobacteriaceae. Clinical Microbiology Reviews, 28(3), 565. 10.1128/CMR.00116-14

Murray, C. J., Ikuta, K. S., Sharara, F., Swetschinski, L., Robles Aguilar, G., Gray, A., Han, C., Bisignano, C., Rao, P., Wool, E., Johnson, S. C., Browne, A. J., Chipeta, M. G., Fell, F., Hackett, S., Haines-Woodhouse, G., Kashef Hamadani, B. H., Kumaran, E. A. P., McManigal, B., … Naghavi, M. (2022). Global burden of bacterial antimicrobial resistance in 2019: a systematic analysis. The Lancet, 399(10325), 629–655. 10.1016/S0140-6736(21)02724-0

Nishii, K., Möller, M., Foster, R. G., Forrest, L. L., Kelso, N., Barber, S., Howard, C., & Hart, M. L. (2023). A high quality, high molecular weight DNA extraction method for PacBio HiFi genome sequencing of recalcitrant plants. Plant Methods, 19(1). 10.1186/S13007-023-01009-X

Nouws, S., Verhaegen, B., Denayer, S., Crombé, F., Piérard, D., Bogaerts, B., Vanneste, K., Marchal, K., Roosens, N. H. C., & De Keersmaecker, S. C. J. (2023). Transforming Shiga toxin-producing Escherichia coli surveillance through whole genome sequencing in food safety practices. Frontiers in Microbiology, 14, 1204630. 10.3389/FMICB.2023.1204630/FULL

Oehler, J. B., Burns, K., Warner, J., & Schmitz, U. (2025). Long-Read Sequencing for the Rapid Response to Infectious Diseases Outbreaks. Current Clinical Microbiology Reports, 12(1), 10. 10.1007/S40588-025-00247-Y

Ofori, B., Twum, S., Yeboah, S. N., Ansah, F., & Sarpong, K. A. N. (2024). Towards the development of cost-effective point-of-care diagnostic tools for poverty-related infectious diseases in sub-Saharan Africa. PeerJ, 12(6), e17198. 10.7717/PEERJ.17198

Ogunleye, A. O., Ghosh, P., Gueye, A. B., Jemilehin, F. O., Okunlade, A. O., Ogunleye, V. O., Kobialka, R. M., Rausch, F., Tanneberger, F., Ajuwape, A. T. P., Sow, O., Ademowo, G. O., Binsker, U., Abd El Wahed, A., Truyen, U., Dieye, Y., & Fall, C. (2025). Nanopore Sequencing-Driven Mapping of Antimicrobial Resistance Genes in Selected Escherichia coli Isolates from Pigs and Poultry Layers in Nigeria. Antibiotics, 14(8), 827. 10.3390/ANTIBIOTICS14080827/S1

Pai, N. P., Vadnais, C., Denkinger, C., Engel, N., & Pai, M. (2012). Point-of-Care Testing for Infectious Diseases: Diversity, Complexity, and Barriers in Low- And Middle-Income Countries. PLoS Medicine, 9(9). 10.1371/JOURNAL.PMED.1001306

Papamentzelopoulou, M., Vrioni, G., & Pitiriga, V. (2025). Comparative Evaluation of Sequencing Technologies for Detecting Antimicrobial Resistance in Bloodstream Infections. Antibiotics 2025, Vol. 14, Page 1257, 14(12), 1257. 10.3390/ANTIBIOTICS14121257

Peker, N., Schuele, L., Kok, N., Terrazos, M., Neuenschwander, S. M., De Beer, J., Akkerman, O., Peter, S., Ramette, A., Merker, M., Niemann, S., Couto, N., Sinha, B., & Rossen, J. W. A. (2021). Evaluation of whole-genome sequence data analysis approaches for short- and long-read sequencing of Mycobacterium tuberculosis. Microbial Genomics, 7(11), 000695. 10.1099/MGEN.0.000695/CITE/REFWORKS

Pheiffer, F., Schneider, Y. K. H., Hansen, E. H., Andersen, J. H., Isaksson, J., Busche, T., Rückert, C., Kalinowski, J., Zyl, L. van, & Trindade, M. (2022). Bioassay-Guided Fractionation Leads to the Detection of Cholic Acid Generated by the Rare Thalassomonas sp. Marine Drugs 2023, Vol*. 21*, 21(1). 10.3390/MD21010002

Pokharel, S., Raut, S., & Adhikari, B. (2019). Tackling antimicrobial resistance in low-income and middle-income countries. BMJ Global Health, 4(6), 2104. 10.1136/BMJGH-2019-002104

Prieto-Fernández, F., Lambert, S., & Kujala, K. (2024). Assessment of microbial communities from cold mine environments and subsequent enrichment, isolation and characterization of putative antimony- or copper-metabolizing microorganisms. Frontiers in Microbiology, 15, 1386120. 10.3389/FMICB.2024.1386120/FULL

Procop, G. W., Nelson, S. K., Blond, B. J., Souers, R. J., & Massie, L. W. (2020). The Impact of Transit Times on the Detection of Bacterial Pathogens in Blood Cultures: A College of American Pathologists Q-Probes Study of 36 Institutions. Archives of Pathology & Laboratory Medicine, 144(5), 564–571. 10.5858/ARPA.2019-0258-CP

Pronyk, P. M., de Alwis, R., Rockett, R., Basile, K., Boucher, Y. F., Pang, V., Sessions, O., Getchell, M., Golubchik, T., Lam, C., Lin, R., Mak, T. M., Marais, B., Twee-Hee Ong, R., Clapham, H. E., Wang, L., Cahyorini, Y., Polotan, F. G. M., Rukminiati, Y., … Sintchenko, V. (2023). Advancing pathogen genomics in resource-limited settings. Cell Genomics, 3(12), 100443. 10.1016/J.XGEN.2023.100443

Purushothaman, S., Meola, M., Roloff, T., Rooney, A. M., & Egli, A. (2024). Evaluation of DNA extraction kits for long-read shotgun metagenomics using Oxford Nanopore sequencing for rapid taxonomic and antimicrobial resistance detection. Scientific Reports 2024 14:1, 14(1), 29531-. 10.1038/s41598-024-80660-3

Roach, D. J., Sangruji, B. P., Bhat, S., Tesfamariam, S., Ben-Zion, I., Bern, M., Bagnall, J., Shoresh, N., Milien, L., & Bhattacharyya, R. P. (2025). BADLOCK: A Rapid, Portable, Inexpensive Diagnostic for Bacterial Pathogen and Resistance Detection in Resource-Limited Settings. MedRxiv, 2025.08.11.25332217. 10.1101/2025.08.11.25332217

Robertson, J., & Nash, J. H. E. (2018). MOB-suite: software tools for clustering, reconstruction and typing of plasmids from draft assemblies. Microbial Genomics, 4(8), e000206. 10.1099/MGEN.0.000206/CITE/REFWORKS

Rozwandowicz, M., Brouwer, M. S. M., Fischer, J., Wagenaar, J. A., Gonzalez-Zorn, B., Guerra, B., Mevius, D. J., & Hordijk, J. (2018). Plasmids carrying antimicrobial resistance genes in Enterobacteriaceae. Journal of Antimicrobial Chemotherapy, 73(5), 1121–1137. 10.1093/JAC/DKX488

*RStudio Desktop - Posit*. (2025). Retrieved December 1, 2025, from https://posit.co/download/rstudio-desktop/

Sakai, J., Tarumoto, N., Kodana, M., Ashikawa, S., Imai, K., Kawamura, T., Ikebuchi, K., Murakami, T., Mitsutake, K., Maeda, T., & Maesaki, S. (2019). An identification protocol for ESBL-producing Gram-negative bacteria bloodstream infections using a MinION nanopore sequencer. Journal of Medical Microbiology, 68(8), 1219–1226. 10.1099/jmm.0.001024

Sanders, E. R. (2012). Aseptic Laboratory Techniques: Plating Methods. J. Vis. Exp, 63, 3064. 10.3791/3064

Schmidt, H., & Hensel, M. (2004). Pathogenicity Islands in Bacterial Pathogenesis. Clinical Microbiology Reviews, 17(1), 14. 10.1128/CMR.17.1.14-56.2004

Schurig, S., Ceruti, A., Wende, A., Lübcke, P., Eger, E., Schaufler, K., Frimpong, M., Truyen, U., Kobialka, R. M., & Abd El Wahed, A. (2024). Rapid Identification of Bacterial Composition in Wastewater by Combining Reverse Purification Nucleic Acid Extraction and Nanopore Sequencing. ACS ES&T Water, 4(4), 1808–1818. 10.1021/ACSESTWATER.3C00794

Schurig, S., Kobialka, R., Wende, A., Ashfaq Khan, M. A., Lübcke, P., Eger, E., Schaufler, K., Daugschies, A., Truyen, U., & Abd El Wahed, A. (2023). Rapid Reverse Purification DNA Extraction Approaches to Identify Microbial Pathogens in Wastewater. Microorganisms, 11(3), 813. 10.3390/MICROORGANISMS11030813/S1

Seale, A. C., Gordon, N. C., Islam, J., Peacock, S. J., & Scott, J. A. G. (2017). AMR surveillance in low and middle-income settings - A roadmap for participation in the Global Antimicrobial Surveillance System (GLASS). Wellcome Open Research, 2. 10.12688/WELLCOMEOPENRES.12527.1

Shen, W., Sipos, B., & Zhao, L. (2024). SeqKit2: A Swiss army knife for sequence and alignment processing. IMeta, 3(3). 10.1002/IMT2.191

Sidstedt, M., Rådström, P., & Hedman, J. (2020). PCR inhibition in qPCR, dPCR and MPS—mechanisms and solutions. Analytical and Bioanalytical Chemistry, 412(9), 2009. 10.1007/S00216-020-02490-2

Siriken, B. (2013). [Salmonella pathogenicity islands]. Mikrobiyoloji Bulteni, 47(1), 181–188. 10.5578/MB.4138

Slattery, S., Tony Pembroke, J., Murnane, J. G., & Ryan, M. P. (2020). Isolation, nucleotide sequencing and genomic comparison of a Novel SXT/R391 ICE mobile genetic element isolated from a municipal wastewater environment. Scientific Reports 2020 10:1, 10(1), 8716-. 10.1038/s41598-020-65216-5

Smith, A. D., & de Sena Brandine, G. (2021). Falco: high-speed FastQC emulation for quality control of sequencing data. F1000Research 2021 8:1874, 8, 1874. 10.12688/f1000research.21142.2

Thornval, N. R., Lacy-Roberts, N., Nilsson, P., Espinosa-Gongora, C., Hasman, H., Mourão, J., Rasmussen, A., Rebelo, A. R., Gibson, C., & Hendriksen, R. S. (2025). Evaluation of four DNA extraction kits for implementation of nanopore sequencing in routine surveillance of antimicrobial resistance in low-resource settings. Frontiers in Microbiology, 16, 1715467. 10.3389/FMICB.2025.1715467/BIBTEX

Timilsina, M., Chundru, D., Pradhan, A. K., Blaustein, R. A., & Ghanem, M. (2025). Benchmarking Metagenomic Pipelines for the Detection of Foodborne Pathogens in Simulated Microbial Communities. Journal of Food Protection, 88(9), 100583. 10.1016/J.JFP.2025.100583

Wasswa, F. B., Kassaza, K., Nielsen, K., & Bazira, J. (2022). MinION Whole-Genome Sequencing in Resource-Limited Settings: Challenges and Opportunities. Current Clinical Microbiology Reports, 9(4), 52. 10.1007/S40588-022-00183-1

Wick, R. R., Volkening, J., & Loman, N. (2017). *Porechop*.

Wood, D. E., & Salzberg, S. L. (2014). Kraken: ultrafast metagenomic sequence classification using exact alignments. Genome Biology 2014 15:3, 15(3), R46-. 10.1186/GB-2014-15-3-R46

World Health Organization. (2014). *Antimicrobial resistance: global report on surveillance*.

Yamin, D., Uskoković, V., Wakil, A. M., Goni, M. D., Shamsuddin, S. H., Mustafa, F. H., Alfouzan, W. A., Alissa, M., Alshengeti, A., Almaghrabi, R. H., Fares, M. A. A., Garout, M., Al Kaabi, N. A., Alshehri, A. A., Ali, H. M., Rabaan, A. A., Aldubisi, F. A., Yean, C. Y., & Yusof, N. Y. (2023). Current and Future Technologies for the Detection of Antibiotic-Resistant Bacteria. Diagnostics, 13(20), 3246. 10.3390/DIAGNOSTICS13203246

Yang, S., & Rothman, R. E. (2004). PCR-based diagnostics for infectious diseases: Uses, limitations, and future applications in acute-care settings. Lancet Infectious Diseases, 4(6), 337–348. 10.1016/S1473-3099(04)01044-8

Yoshida, C. E., Kruczkiewicz, P., Laing, C. R., Lingohr, E. J., Gannon, V. P. J., Nash, J. H. E., & Taboada, E. N. (2016). The Salmonella In Silico Typing Resource (SISTR): An Open Web-Accessible Tool for Rapidly Typing and Subtyping Draft Salmonella Genome Assemblies. PLOS ONE, 11(1), e0147101. 10.1371/JOURNAL.PONE.0147101

Zhang, S., den Bakker, H. C., Li, S., Chen, J., Dinsmore, B. A., Lane, C., Lauer, A. C., Fields, P. I., & Deng, X. (2019). SeqSero2: Rapid and improved salmonella serotype determination using whole-genome sequencing data. Applied and Environmental Microbiology, 85(23). 10.1128/AEM.01746-19

Zhou, Z., Alikhan, N.-F., Mohamed, K., Fan, Y., Group, the A. S., Achtman, M., Brown, D., Chattaway, M., Dallman, T., Delahay, R., Kornschober, C., Pietzka, A., Malorny, B., Petrovska, L., Davies, R., Robertson, A., Tyne, W., Weill, F.-X., Accou-Demartin, M., & Williams, N. (2020). The EnteroBase user’s guide, with case studies on Salmonella transmissions, Yersinia pestis phylogeny, and Escherichia core genomic diversity. Genome Research, 30(1), 138–152. 10.1101/GR.251678.119

